# Unique Hmgn2 Orthologous Variant Modulates Shape Preference Behavior in Medaka Fish

**DOI:** 10.1101/2023.11.17.567541

**Authors:** Yume Masaki, Shuntaro Inoue, Shinichi Nakagawa, Saori Yokoi

**Author notes:** Correspondence should be addressed to Saori Yokoi.

## Abstract

Diversification of protein sequences contributes to the variation of physiological traits. In this study, using medaka fish (*Oryzias latipes*) as a model, we identified a novel protein variant influencing shape preference behavior. Re-analysis of sequencing data revealed that *LOC101156433*, previously annotated as a non-protein-coding gene, encodes a divergent Hmgn2 variant undetectable by standard homology searches. This unique Hmgn2 in medaka shows non-standard subnuclear localization, and mutants exhibited a reduction in certain regions of the telencephalon and a loss of shape preference. These results not only establish a direct association between amino acid sequence variation and the development of new molecular properties and behavioral adaptations but also highlight new clues toward understanding the visual shape perception system in fish.

## Introduction

The molecular basis for the diversity of species remains an enigma, given the relative consistency in the number of protein-coding genes observed across multicellular organisms, especially across vertebrate species. For instance, the human genome comprises approximately 20,000 protein-coding genes, a number similar to that in other vertebrate species, including mouse, chicken, lizard, frog, and fish^1,2^. The central question is how organisms with comparable numbers of protein-coding genes create various morphological and physiological traits. A prevailing theory in molecular biology is that diversification primarily arises from modifications in cis-regulatory elements including enhancers, silencers, and insulators^3,4^. Changes in these regulatory domains can lead to differential gene expression patterns, even in the context of conserved protein-coding sequences, contributing to phenotypic variation^5^.

Beyond cis-regulatory changes, the emergence of novel proteins with distinct amino acid sequences might also play a pivotal role in evolutionary processes. Indeed, recent studies have revealed that a number of species-specific short peptides are produced from transcripts once annotated as long noncoding RNAs, which introduce new functionalities that can enable organisms to acquire unique traits and adapt to varied ecological niches^6,7^. In addition, it has been proposed that the introduction of particular mutations in certain proteins dramatically changes their biochemical properties, enabling behavioral changes that lead to adaptation to specific environments^8^. These perspectives underscore the importance of considering both regulatory and protein-coding adaptations in our understanding of molecular evolution.

Shape perception, a cognitive function essential for the survival of many species, enables organisms to navigate complex environments, identify resources, escape predators, and engage in social interactions^9,10^. The role of shape perception goes beyond mere visual processing; it plays a key part in learning, memory, and decision-making^11,12^. Thus, the proficiency of an organism in discerning shapes can have profound implications on its adaptive strategies and overall survival^13^. The understanding of shape perception mechanisms is well-established in mammalian species, yet remains to be fully explored in non-mammalian species.

Medaka fish (*Oryzias latipes*) provides an advantageous model for probing the genetic basis of behavior, given its established genetic resources, including CRISPR/Cas9^14,15^, and its sophisticated visual system that is vital for behaviors like individual recognition and object discrimination^16,17,18,19^.

This study utilizes the medaka’s genetic tractability to investigate new genetic elements that influence behavior, focusing on short peptides derived from genes initially annotated as lncRNAs. Our research uncovered a medaka ortholog of the Hmgn2 gene, which displays such a sequence divergence that conventional homology searches failed to identify it. Notably, this medaka Hmgn2 homolog exhibits unique subnuclear localizations, and *Hmgn2* mutant medaka exhibited changes in shape preference. These findings point to the essential role of amino acid diversification in medaka, impacting not merely molecular characteristics but also significantly influencing the capability for shape perception.

## Results

### Identification of putative protein-coding lncRNAs in the adult medaka brain

Following a reanalysis of RNA-seq and Ribo-seq from adult medaka brain samples^20^, we closely examined 3,096 genes annotated as lncRNAs (NCBI Annotation Release 103) for evidence of translation. As a result, RNA-seq analysis indicated that 743 of these genes were expressed (TPM > 2) in the adult brain. Further examination of translation efficiency, determined by the ratio of TPM^Ribo^ to TPM^RNA^ (TPM^Ribo^ / TPM^RNA^), revealed that 124 out of these 743 genes had a translation efficiency of 0.5 or higher, suggestive of active translation (Fig. 1a, Supplementary Dataset 1). Two examples of lncRNA-encoded proteins are shown in Fig. 1b. Ribo-seq reads specifically mapped to exonic regions of *LOC111946391* and *LOC101156433*, implying these sites as active translation regions (Fig. 1b). The genes with the highest number of reads from both RNA-seq and Ribo-seq are listed in Table 1. BLAST searches for amino acid sequences derived from Ribo-seq results identified potential homologous proteins, with LOC111946391 and LOC101159365 corresponding to tmsb4 and DSS1, respectively. In contrast, LOC101156433 did not match any known protein homologs, leading us to further investigate its potential as a novel protein-coding gene.

**Fig. 1.**
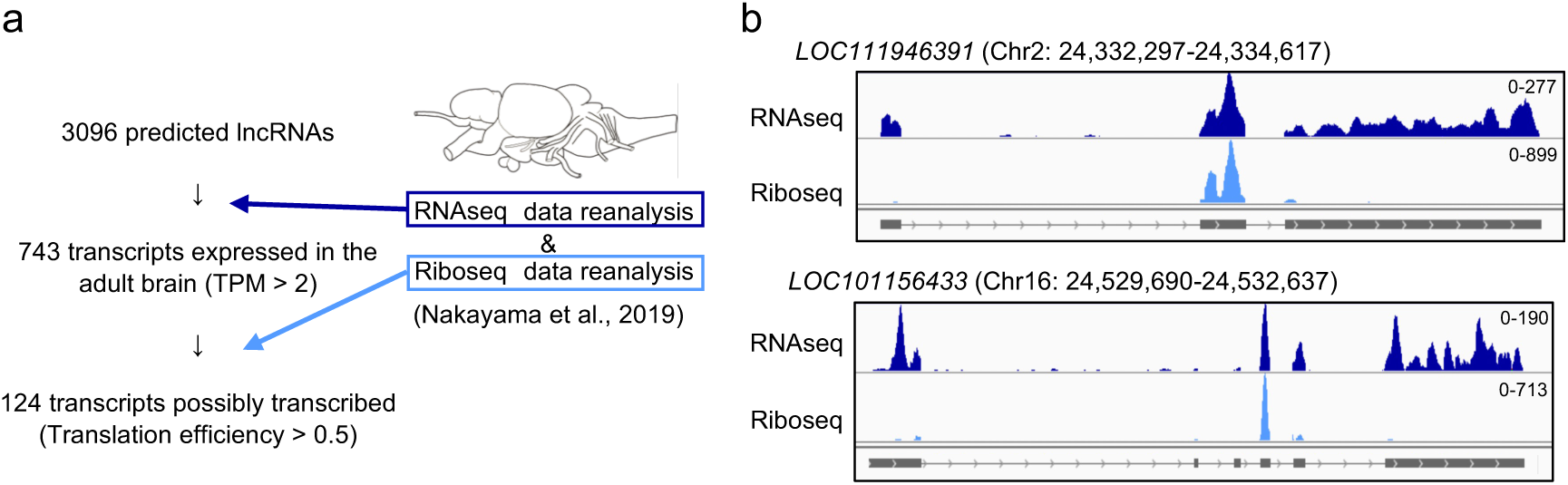
Identification of coding genes annotated as lncRNAs. a. Overview of public data reanalysis. b. Coverage tracks for RNA-Seq (dark blue) and Ribo-seq (light blue) of *LOC111946391* and *LOC101156433*, identified as potentially translated. The tracks show the distribution and density of sequencing reads along the length of these genes. Peaks in the Ribo-Seq data, relative to RNA-Seq, suggest active translation in regions previously annotated as non-coding, indicating these genes may in fact encode proteins.

**Table. 1.** The top 5 highly expressed genes predicted to be protein-coding lncRNA through RNA-seq and Ribo-seq reanalysis. This table lists the top five genes that were highly expressed in the RNA-seq reanalysis and are predicted to be protein-coding lncRNAs. The table includes information on each gene’s expression levels in both RNA- seq and Ribo-seq data, alongside their predicted peptide homology. Out of these five genes, four were identified as potential homologs of known peptides, suggesting a coding potential previously unrecognized in these lncRNA annotations.

| Gene name | TPM<br>(RNAseq) | Translation<br>efficiency | BLAST top hit protein |
| --- | --- | --- | --- |
| *LOC111946391* | 365.476384 | 2.21365351 | Tmsb4 (*Cyprinodon variegatus*)<br>(E value = 8e-18) |
| *LOC101159365* | 355.257276 | 1.30817715 | DSS1 (*Liparis tanakae*)<br>(E value = 8e-42) |
| *LOC101156433* | 327.5903 | 0.90638175 | Hypothetical protein FQA47_008997(*Oryzias melastigma*)<br>(E value = 6e-33) |
| *LOC101155702* | 288.781364 | 2.02049597 | Atp5me (*Perca fluviatilis*)<br>(E value = 2e-36) |
| *LOC111948018* | 256.747376 | 0.79612099 | Tmsb1 (*Neolamprologus brichardi*)<br>(E value = 4e-16) |

### LOC101156433 is a divergent homolog of Hmgn2

As the annotation of *LOC101156433* was predicted by automated computational analysis using Gnomon, we performed 5’ and 3’ rapid amplification of cDNA ends (RACE) and obtained the full-length cDNA for *LOC101156433*. The sequence of *LOC101156433* cDNA we obtained (deposited in GenBank with accession number LC782854) is 99-bp shorter in the 5’UTR compared to the public annotation but otherwise consistent across other regions. Therefore, we excluded the annotation error as the reason for the absence of homology in BLAST searches. Interpro domain search^21^ predicted that LOC101156433 belonged to the Hmgn family, and synteny comparisons suggested it is an ortholog of Hmgn2 (Fig. 2a), henceforth referred to as oHmgn2 (Hmgn2 in *oryzias latipes*). HMGN proteins are non-histone proteins that play crucial roles in chromatin remodeling and gene expression regulation^22^. They are characterized by a conserved domain architecture, including a bipartite nuclear localization signal (NLS), a nucleosome-binding domain (NBD), and a regulatory domain^22^. These domains are essential for the proteins’ interaction with nucleosomes and their subsequent function in chromatin dynamics^23,24^.

**Fig. 2.**
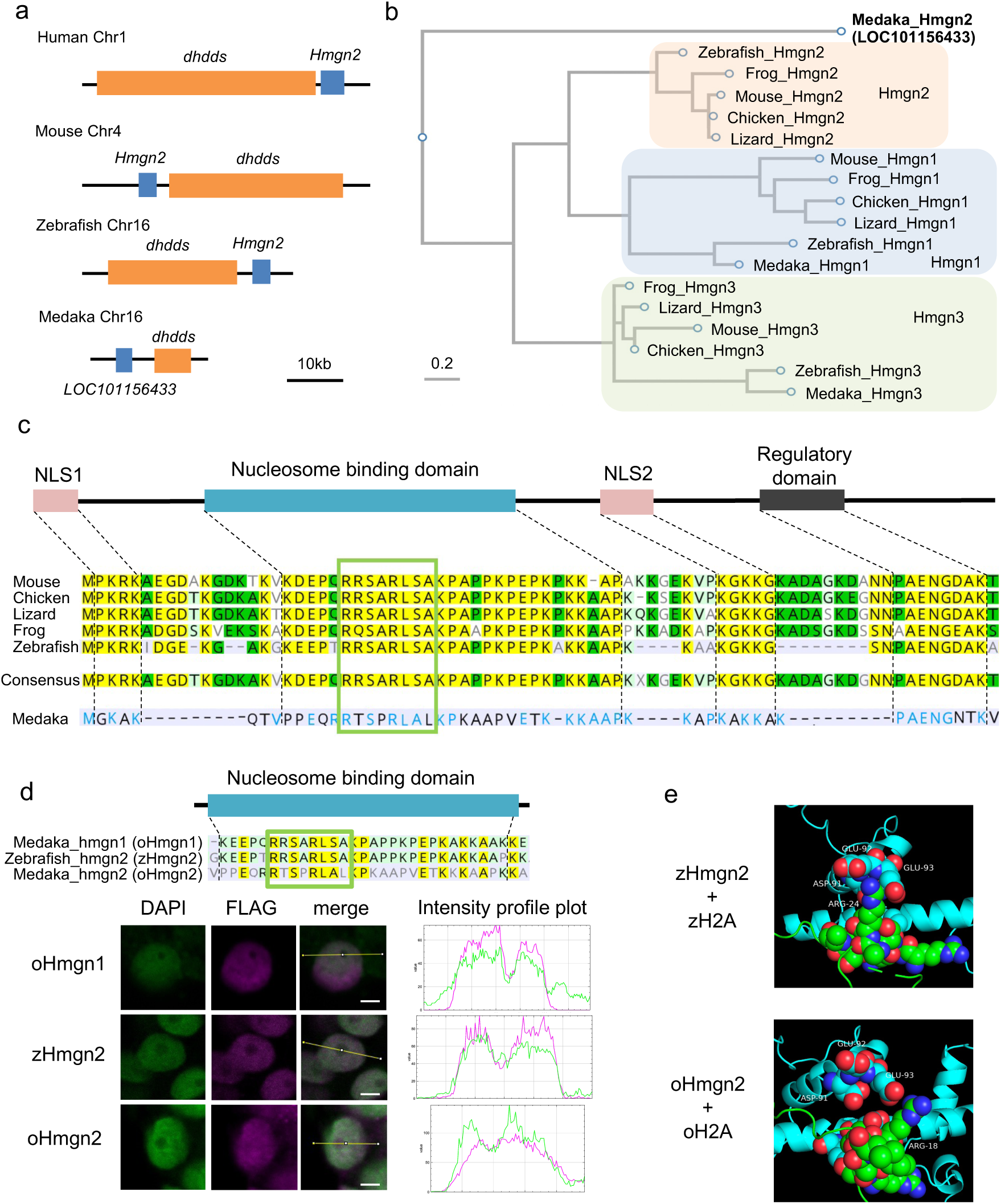
Characterization of *LOC101156433* (*oHmgn2*). a. The genomic loci of *dhdds* and *Hmgn2* in humans, mice, and zebrafish, alongside the orthologous region in medaka. b. A maximum likelihood (ML) phylogenetic tree illustrating the relationships among Hmgn1, Hmgn2, and Hmgn3 protein lineages from various species. Branch supports are indicated by Chi2-based parametric values from the approximate likelihood ratio test. Notably, LOC101156433 does not cluster within any established Hmgn clades, suggesting its unique evolutionary trajectory. c. An alignment of Hmgn2 protein sequences from representative vertebrates, including medaka, is shown. The consensus sequence (representing mouse, chicken, lizard, frog, and zebrafish) is compared with medaka’s sequence. Amino acid similarities are color-coded: yellow for 100% similarity, green for 80-100%, light green for 60-80%, and unhighlighted for less than 60%. Similarities between the consensus sequence and medaka’s sequence are highlighted in blue. The nucleosome binding core domain is marked by an open green box. d. The protein sequence alignment for oHmgn1, zHmgn2, and oHmgn2, focusing on their nucleosome binding core domains (open green box). oHmgn1 and zHmgn2 possess the complete domain “RRSARLSA”. Intensity profile plots generated along the indicated yellow lines in the left images reveal different staining patterns; oHmgn1 and zHmgn2 align with DAPI staining, suggesting nucleoplasm localization, while oHmgn2 does not. Scale bars measure 2 μm. e. The predicted interactions between H2A (light blue) and Hmgn2 (green) in zebrafish and medaka, using AlphaFold2. The nucleosome binding core domain “RRSARLSA” in Hmgn2 and key positively charged residues in H2A required for binding are shown as spheres. The color coding for the atomic structures is as follows: oxygen atoms in red and nitrogen atoms in blue. The model predicts a specific interaction in zebrafish, where the side chain of R24 in zHmgn2 appears to engage with the D91-E93 residues of H2A. In contrast, such an interaction is not predicted for oHmgn2 in medaka, highlighting a potential structural and functional divergence between the species.

### Distinct characteristics and evolutionary divergence of oHmgn2

Next, a phylogenetic tree was constructed using the amino acid sequences of Hmgn1, Hmgn2, and Hmgn3 in various vertebrates. Consequently, while medaka Hmgn1 and Hmgn3 aligned with their respective clades, oHmgn2 clustered separately from the Hmgn2 clades of other species (Fig. 2b). This phylogenetic distinction underscored oHmgn2’s unique characteristics as a divergent ortholog within the HMGN protein family. Sequence analysis of the HMGN2 protein across five vertebrate species (mouse, chicken, lizard, frog, and zebrafish) revealed high homology within the four key domains: NLS1, NBD, NLS2, and the regulatory domain. In contrast, oHmgn2 from medaka displayed significantly lower conservation in these domains. The most striking variation was found in the NBD, where the core sequence ‘RRSARLSA’, critical for nucleosome interaction^23^, was completely conserved in the other five species but only 60% conserved in medaka (Fig. 2c). Previous research has demonstrated that alterations in this core sequence could mislocalize HMGN to the nucleolus and reduce its affinity for chromatin^24^.

To ascertain the subcellular localization of HMGN proteins within medaka cells, we compared their intranuclear positioning against medaka Hmgn1 (oHmgn1) and zebrafish Hmgn2 (zHmgn2), both of which contain the intact core sequence. The coding sequences of each HMGN gene were fused with a FLAG tag sequence and transcribed. These mRNA constructs were then microinjected into the one-cell stage of medaka embryos. After a 24-hour incubation period, the spatial distribution of the HMGN proteins was detected by immunostaining with an anti-FLAG antibody (Supplementary Figure 1). DAPI staining typically produces a less intense signal in the nucleolus due to its lower DNA content compared to the nucleoplasm^25^. In our study, both oHmgn1 and zHmgn2 showed localization patterns that mirrored DAPI staining, suggesting their presence predominantly in the nucleoplasm rather than the nucleolus (Fig. 2d). On the other hand, oHmgn2 demonstrated a broader localization pattern (Fig. 2d). While it was clearly present within the nucleus, as indicated by co-localization with DAPI-stained regions, oHmgn2 also appeared in areas of weaker DAPI signal. This implied that oHmgn2 was not exclusively confined to the nucleoplasm but was also present in the nucleolus, differentiating its intranuclear distribution from that of the other HMGN proteins analyzed (Fig. 2d). This differential localization pattern of oHmgn2 could reflect distinct functional roles or interactions specific to this medaka protein variant.

Previous studies have indicated that the function of HMGN2 involves binding to the histone H2A, with specific positively charged residues being essential for this interaction^23^. Utilizing AlphaFold2^26,27^, we modeled the conformation of HMGN2 and H2A to predict their interaction within the nucleosome (Supplementary Movies 1-2). The zebrafish Hmgn2 (zHmgn2) nucleosome binding core domain (RRSARLSA) was predicted to be closely interacting with H2A, specifically, the side chain of R24 possibly engaging with the D91-E93 residues of H2A, which aligned with previous NMR findings^23^ (Fig. 2e). However, for oHmgn2 in medaka, the interaction between its R18 residue and the D91-E93 residues of H2A appeared less pronounced. This suggested a potentially weaker interaction of oHmgn2 with nucleosomes compared to the typical HMGN2-nucleosome interactions observed in other species, pointing to a diminished chromatin-regulatory capacity in oHmgn2.

Extensive phylogenetic analysis through TBLASTN to locate homologs of zHmgn2 and oHmgn2 in ray-finned fishes disclosed distinctive patterns of occurrence (Supplementary Figure 2). In non-Acanthopterygii groups, homologs resembling zHmgn2 were present, while medaka-like oHmgn2 homologs were absent. In contrast, within Acanthopterygii fishes—except those belonging to the Tetraodontiformes order— oHmgn2 homologs were prevalent, and zHmgn2 counterparts were not found. These findings suggested that a significant divergence in the amino acid sequences of Hmgn2 might have occurred within the Acanthopterygii lineage, potentially contributing to the acquisition of novel and distinct properties.

### Expression patterns of oHmgn2 in adult medaka brain neurogenic regions

In line with observations in mammalian systems, specifically the adult mouse brain where Hmgn2 is predominantly expressed in neural progenitor cells^28^, we investigated the expression profile of *oHmgn2* in the adult medaka brain. Notably, *oHmgn2* showed localized expression in neurogenic regions^29,30^, spanning from the optic tectum to the telencephalon (Fig. 3a). This expression pattern was further characterized by the co-expression of the proliferation marker PCNA and the intermediate precursor marker gene *eomesa* (Figs. 3b-c), consistent with an active role in neurogenesis. This finding indicated that oHmgn2, akin to its mammalian homologs, was involved in neural progenitor cell function within the medaka brain.

**Fig. 3.**
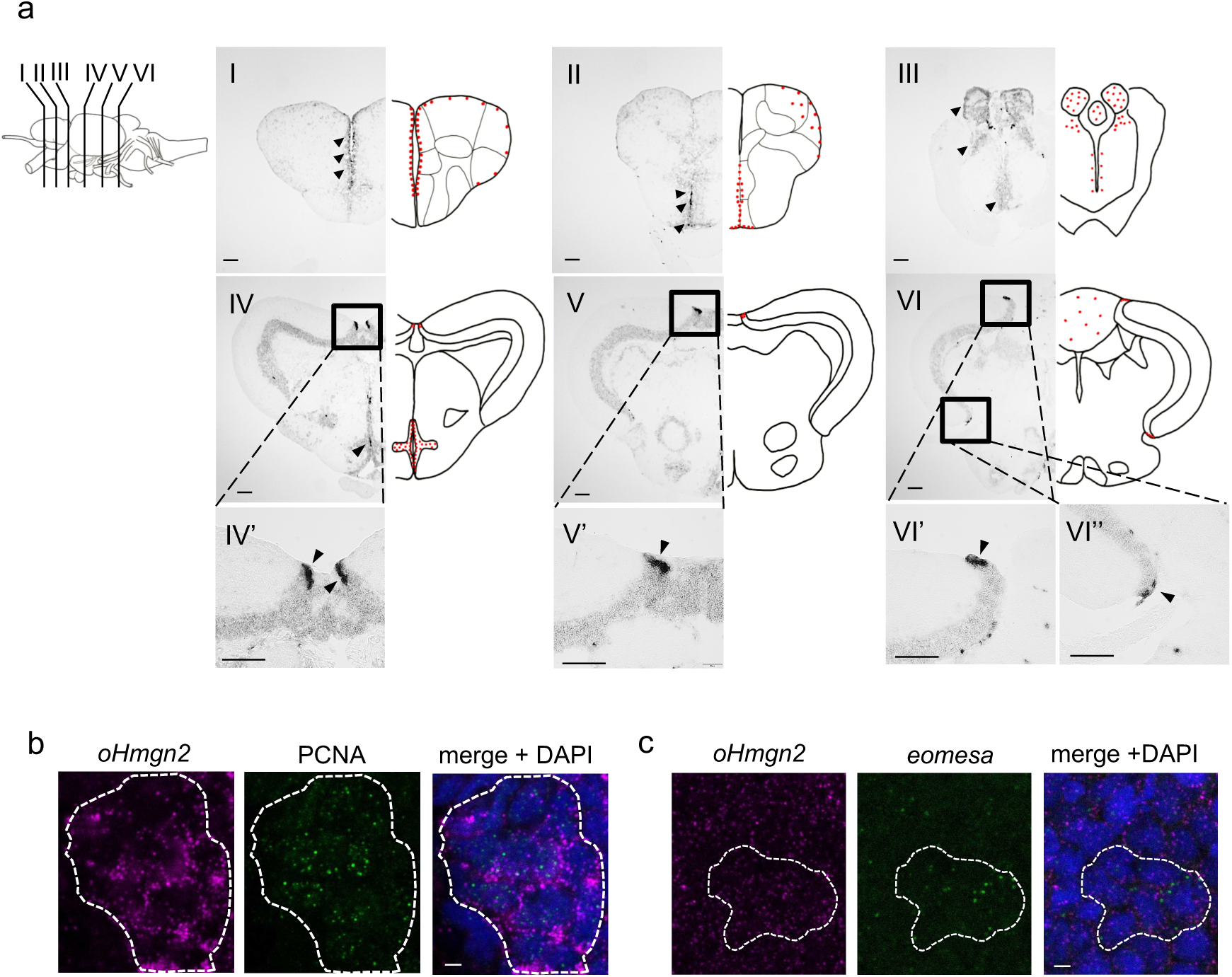
Expression pattern of *oHmgn2* in the adult medaka brain. a. Lateral view of the adult medaka brain (upper left), detailing the positions of six coronal sections (I-VI) indicated by lines. Red dots mark the locations of proliferating cells, as previously reported in Kuroyanagi *et al.*, 2010^29^. Each scale bar in the overview represents 100 μm. b. A representative micrograph displaying cells that co-express *oHmgn2* and PCNA in the tectal marginal zone (referenced in Fig. 3a-IV’, indicated with an arrowhead). A scale bar represents 2 μm. c. A representative micrograph displaying cells that co-express *oHmgn2* and *eomesa*, located in the medial zones of the telencephalon (as shown in Fig. 3a-I, marked with an arrowhead). A scale bar represents 2 μm.

### Viability and morphological assessment of oHmgn2 knockout medaka

To investigate the physiological function of oHmgn2, we generated knockout medaka lines using CRISPR/Cas9, leading to a 2-bp deletion in the third exon. This mutation resulted in a frameshift and subsequent loss of the regulatory domain, suggesting a loss-of-function mutation in these medaka (Fig. 4a, Supplementary Figure 3). Homozygous mutants for this allele were viable and were produced at expected Mendelian ratios (Fig. 4b), with body lengths comparable to wild-type (WT) counterparts (Figs. 4c-d). Brain size measurements between *oHmgn2* homozygous mutants and WT fish showed no significant overall differences (Figs. 4e-f and Supplementary Figure 4), indicating that the loss of *oHmgn2* did not affect overall brain development or fish growth.

**Fig. 4.**
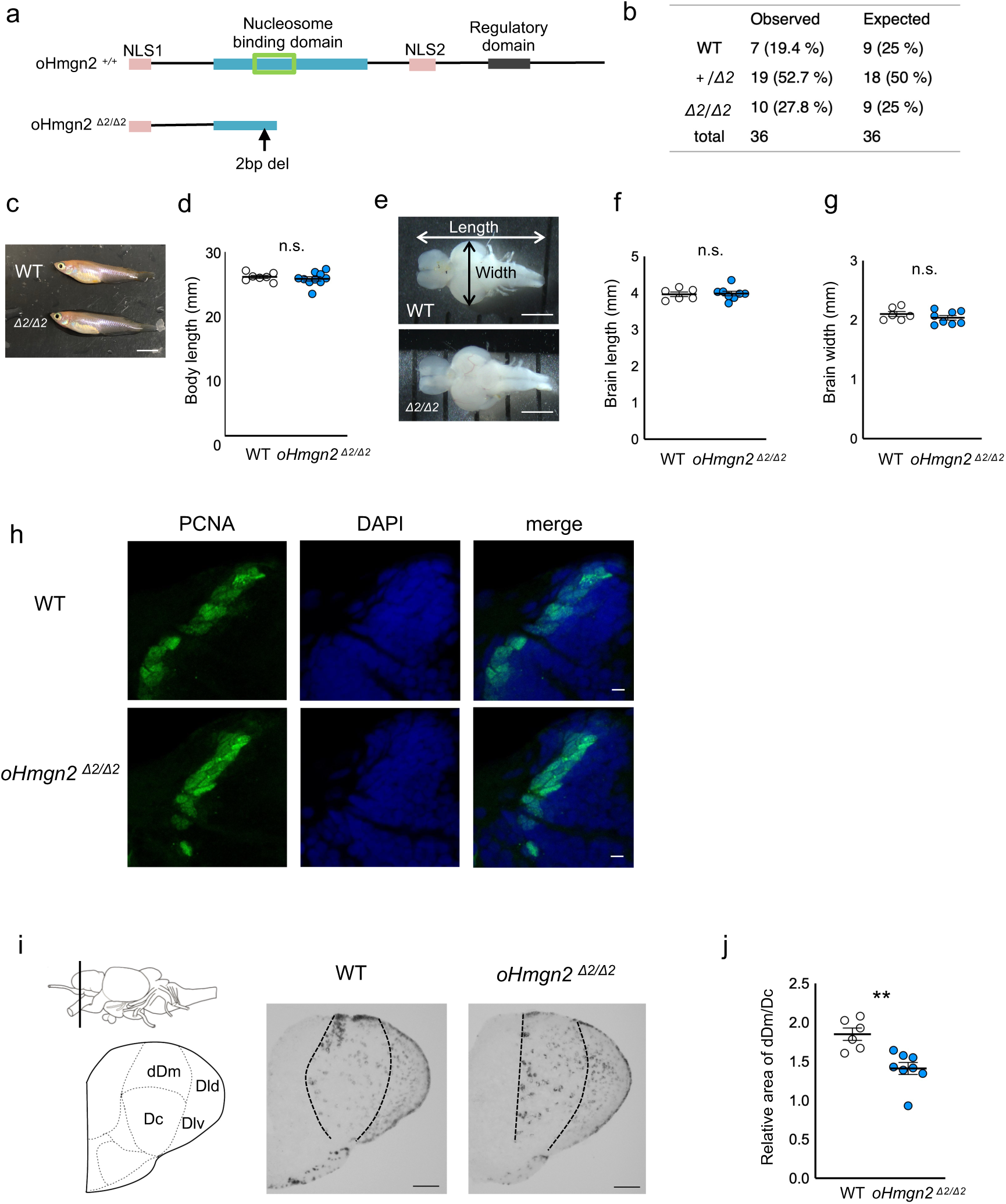
Viability and brain development of *oHmgn2* mutant medaka. a. Outline of *oHmgn2* mutant (*oHmgn2 ^Δ^*^2^*^/Δ^*^2^). In this mutant, the oHmgn2 gene is likely truncated, resulting in the loss of a regulatory domain. b. Table summarizing the genotype distribution of adult fish bred from heterozygous mutants, alongside the expected Mendelian distribution. This data provides insight into the viability of the *oHmgn2* mutants. c. A photograph comparing the body shapes of WT and *oHmgn2* mutant medaka at 5 months post-hatching. A scale bar of 5 mm is provided for size reference. d. Body length of WT and *oHmgn2* mutants. WT, n = 7; *oHmgn2 ^Δ^*^2^*^/Δ^*^2^, n = 10. e. Photographs of the brains of WT and *oHmgn2* mutants. A scale bar represents 1 mm. f. Brain length of WT and *oHmgn2* mutants. WT, n = 6; *oHmgn2 ^Δ^*^2^*^/Δ^*^2^, n = 8. g. Brain width of WT and *oHmgn2* mutants. WT, n = 6; *oHmgn2 ^Δ^*^2^*^/Δ^*^2^, n = 8. h. PCNA expression in the tectal marginal zone (referenced in Fig. 3a-IV’) in WT and *oHmgn2* mutants. i. Micrographs illustrating *eomesa* expression in WT and *oHmgn2* mutants. The position of the coronal section is indicated in the upper left. dDm, the dorsal part of dorsomedial telencephalon; Dc, the dorsocentral telencephalon; Dld, the dorsal part of dorsolateral telencephalon; Dlv, the ventral part of dorsolateral telencephalon. Scale bars represent 100 μm. j. Quantification of relative size of the dDm/Dc regions compared to the Dld/Dlv regions in WT and *oHmgn2* mutants. WT, n = 4; *oHmgn2 ^Δ^*^2^*^/Δ^*^2^, n = 6. d, f, g, j. Mean ± SEM, Mann– Whitney U test: *:P<0.05, n.s., not significant.

High expression of *oHmgn2* mRNA was noted in the internal tectal marginal zone, a region of active neurogenesis, where it was co-expressed with the cell proliferation marker PCNA (Fig. 3b). Immunohistochemical analysis showed no significant difference in PCNA-positive cell populations between WT and *oHmgn2* mutants (Fig. 4g), suggesting that the *oHmgn2* mutation did not affect the proliferation rate within this brain region. Further, co-expression of *oHmgn2* mRNA with the transcription factor *eomesa* (Fig. 3c), a marker for intermediate neural progenitors, was observed. In mammals, an increase in ectopic eomes-positive cells has been documented in the ventricular-subventricular zones of *Hmgn2* knockout mice^28^. In addition, *eomesa* serves as a marker for pallium subdivision in zebrafish^31^. *In situ* hybridization in *oHmgn2* mutants indicated that the expression pattern of *eomesa* remained largely unchanged when compared to WT medaka (Supplementary Figure 5), suggesting that the fundamental patterning of neural progenitor cells was intact despite the *oHmgn2* mutation. However, discrepancies were noted in the size of certain brain regions, specifically the dorsomedial (dDm) and dorsocentral (Dc) areas of the telencephalon, which were significantly reduced in *oHmgn2* mutants relative to WT fish (Figs. 4h-j, P = 0.0013). This size reduction was confirmed through DAPI staining, which demarcates nuclear material and thereby outlines the architecture of brain tissue (Supplementary Figure 6).

Taken together, our data suggested that Hmgn2 played a role in the normal development of the telencephalon in medaka.

### Behavioral impacts of oHmgn2 deficiency on shape perception

Our behavioral analysis explored the effects of the *oHmgn2* mutation on medaka fish behavior. Initial assessments confirmed that the *oHmgn2* mutation did not impair locomotor activity or basic visual responses, as evidenced by unaltered performance in standard tests (Supplementary Figure 7). This indicated that the mutation does not affect the general mobility or visual processing of the fish.

We then focused on the role of oHmgn2 in shape perception. Certain brain regions in fish, such as Dm and Dc, are posited to be analogous to the amygdala and cortex in mammals, respectively^32–34^, which are implicated in cognitive processes including shape perception. The specific function of these regions in medaka, however, has not been characterized. Notably, in zebrafish, regions expressing *eomesa*—a gene pattern similar to Dm and Dc in medaka—are known to be activated during shape perception tasks^35^. While this did not necessarily establish homology, it suggested a potential functional similarity.

In our experimental design, a Y-maze test was used, with one arm lined with black circles and the other with black triangles, to determine if medaka exhibit a natural preference for shape (Fig. 5a). WT medaka demonstrated a significant preference for the triangle zone over the circle zone, indicative of inherent shape preference behavior (P < 0.0001). In contrast, *oHmgn2* mutants showed no significant preference, spending equal time in both arms of the maze (Fig. 5b). The “preference index” was calculated to quantitatively measure this behavior, determined by the difference in duration spent in each zone divided by the total duration spent in both zones (Fig. 5a). The results showed that while WT fish had a measurable preference (P = 0.0049), *oHmgn2* mutants exhibited a negligible preference index, further substantiating the absence of shape preference in these mutants (Fig. 5c). Control observations confirmed that there were no differences in total residence time in the zones or total swimming distance between WT and *oHmgn2* mutants (Figs. 5d-e), suggesting that the lack of shape preference was not attributed to alterations in overall activity or locomotion abilities.

**Fig. 5.**
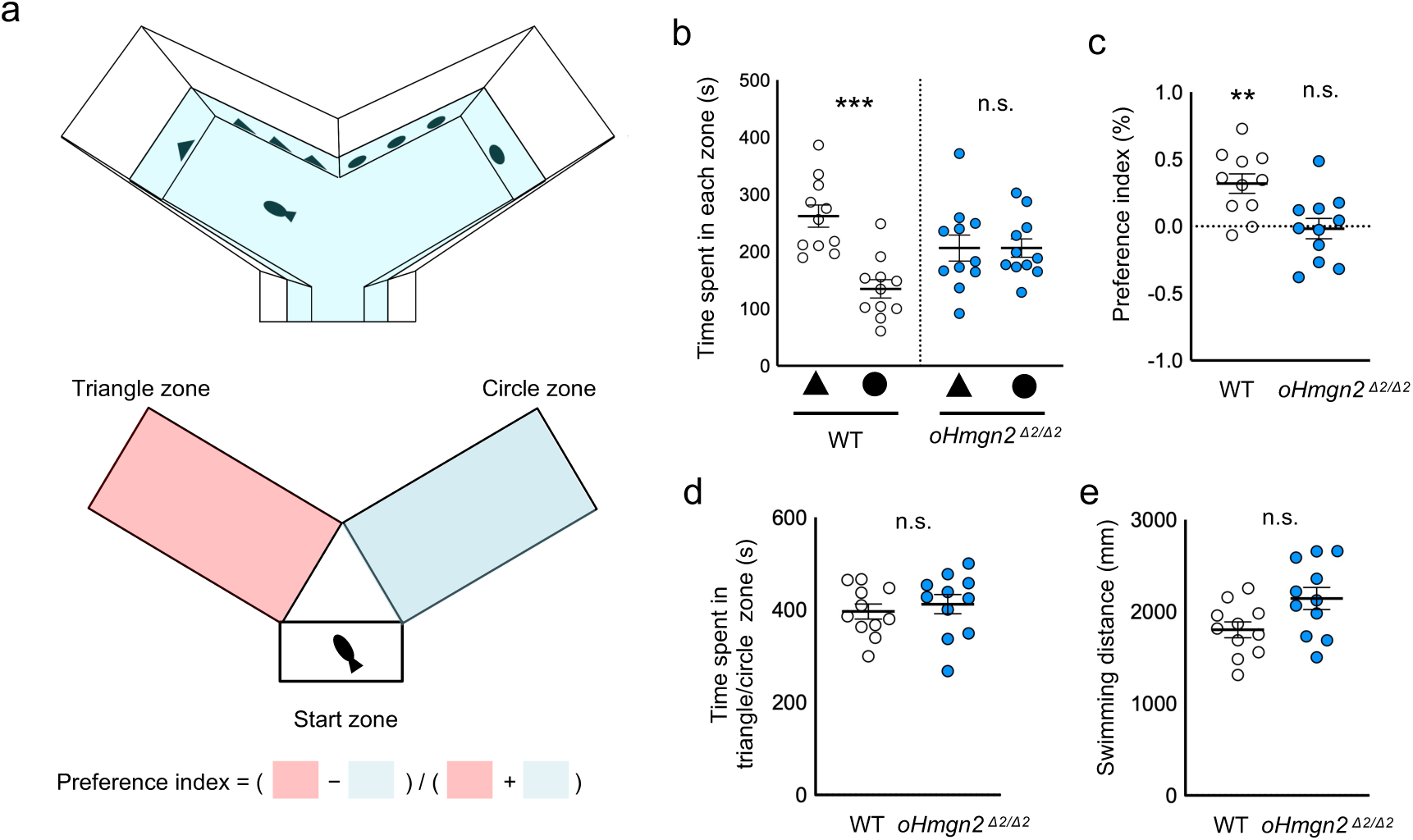
Shape preference of *oHmgn2* mutant medaka. a. The apparatus used for the shape preference test. The Y-Maze apparatus featured two long arms, each with black triangular and circular forms attached to the walls. The test measured the time spent by the fish in the triangle and circle zones to calculate a preference index. b. WT fish spent significantly more time in the triangle zone compared to the circle zone, whereas *oHmgn2* mutants exhibited no such preference. c. WT fish showed a significant preference for the triangular zone, in contrast to *oHmgn2* mutants, which exhibited no preference. One-sample Wilcoxon signed-rank test against a theoretical median of zero. *:P<0.05, n.s., not significant. d. There was no significant difference between WT and *oHmgn2* mutants in the total residence time in either zone. e. There was no significant difference between WT and *oHmgn2* mutants in total swimming distance. b-e. Mean ± SEM, n = 11 per group. b, d, e. Mann– Whitney U test: ***:P<0.001, n.s., not significant.

In summary, *oHmgn2* mutants, despite normal physical and basic sensory motor abilities, demonstrated a deficit in shape perception. This selective behavioral alteration highlights the potential involvement of oHmgn2 in the neural circuit or pathways underlying cognitive functions related to shape perception in medaka fish.

## Discussion

Our comprehensive analysis began with the exploration of lncRNA-annotated genes within the adult medaka brain, leading to the identification of 124 genes with significant translation efficiency, suggesting active translation of these previously thought non-coding regions. The focus on *LOC101156433*, now recognized as a homolog of *Hmgn2* by synteny comparisons, revealed its expression in neural progenitor cells, echoing the expression patterns found in mice. This cross-species consistency in gene expression underscores the evolutionary conservation of HMGN2’s role in neurodevelopment. However, a remarkable divergence in amino acid sequence was observed for oHmgn2 compared to vertebrate counterparts, leading to its nucleolar mislocalization. This unexpected localization, typically associated with dysfunctional HMGN proteins, may indicate a species-specific adaptation in medaka, wherein oHmgn2 performs distinct cellular functions possibly related to the unique attributes of medaka neurobiology. A previous study has indicated that the nucleolar localization of mutated HMGN proteins corresponds with their failure to bind chromatin effectively^36^. The potential weak affinity of oHmgn2 for chromatin could denote a divergent evolutionary pathway, a hypothesis supported by AlphaFold2 structural predictions. Yet, the precise nature of this relationship requires further study to elucidate the chromatin dynamics involving oHmgn2.

In medaka, *oHmgn2* mutants exhibited viability and normal brain size, in contrast to the subviability and microcephaly observed in *Hmgn2* mutant mice. This suggests a divergent role for HMGN2 between species, potentially due to compensatory genetic mechanisms unique to medaka. Notably, the distinct phenotypic presentation in medaka points to a more refined role of oHmgn2 in the development and patterning of specific brain regions rather than a global impact on brain size or cell proliferation. The discrepancy between the mouse and medaka phenotypes raises the question of whether the divergent sequences in medaka confer new functions upon oHmgn2, akin to the proposed adaptive significance of amino acid substitutions in the human FOXP2 gene^8^. Alternatively, the differences could be attributed to a less critical role of oHmgn2 in medaka, possibly as a result of its attenuated interaction with chromatin. Delineating the causes of these phenotypic variations will require in-depth molecular and genetic analyses, potentially involving the assessment of oHmgn2’s interactions with other nuclear components and its broader regulatory influence on gene expression.

The viability of *oHmgn2* mutants in medaka allowed for the unique opportunity to investigate behavioral implications of the gene’s disruption. Our observations revealed a natural preference in WT medaka for triangular shapes over circular ones, a behavioral trait not previously characterized as previous studies have typically focused on more complex, animal-like shapes^37^. While zebrafish are known to avoid certain simple shape^12^, suggesting a shared behavioral trait among teleosts, the underlying reasons for this preference remain to be fully understood. Although it is not possible to determine whether medaka avoided circles or preferred triangles in this experiment alone, one hypothesis is that the avoidance of circular shapes could be an innate response to predator mimicry. This theory is supported by the startle response elicited by rapidly expanding circles, which may simulate an approaching threat^38^, and attack cone avoidance behavior, where they steer clear of shapes that could represent the eyes or mouths of predators^39,40^.

Understanding the neurobiological basis of shape perception in vertebrates, particularly within the ‘what pathway’ of the ventral visual stream^41^, has predominantly been guided by studies in primates and, to a lesser extent, rodents. These studies have revealed specialized brain regions capable of discriminating shapes, such as the lateromedial region of the primary visual cortex in mice^42^, which are thought to contribute to the ‘what pathway’ implicated in object recognition^43^. In teleosts, however, the process of identifying and functionally validating brain regions analogous to those involved in shape perception has been slower to develop, partly due to experimental constraints and the scarcity of appropriate genetic models. The medaka fish, as demonstrated in our study, presents a fresh perspective into this domain of cognitive neuroscience. The *oHmgn2* mutant medaka, which showed no preference for triangles over circles, provides evidence for potential abnormalities in shape perception, suggesting the functional involvement of the dDm and Dc brain regions in these processes.

Our findings in medaka offer intriguing parallels to studies conducted on goldfish and carp^44^. These studies have highlighted that the dorsal zone of the rostrolateral region of the lateral preglomerular nucleus (PGlr-d) – a brain region receiving both direct retinal inputs and indirect inputs via the tectum – extends fibers to the part of dDm and Dc regions. These evolutionarily conserved structures hint at a fundamental mechanism in vertebrate brain organization that underlies visual information processing. However, the evolutionary divergence between the Acanthomorpha group, to which medaka belongs, and the Ostariophysi group, which includes goldfish and carp, introduces complexities in direct functional comparisons^45^. Adding to the complexity, the teleost research field lacks comprehensive functional validation for these specific brain regions. Furthermore, recent research proposes that medaka’s dDm region is compartmentalized into discrete clonal units, each with a unique gene expression signature^46^. Consequently, to clarify the roles of the dDm and Dc regions in medaka, it is crucial to conduct detailed anatomical studies to map out the neural connections and to evaluate the functions of these individual clonal units. Undertaking this task, especially in *oHmgn2* mutant medaka, holds the potential to illuminate the underpinnings of shape perception in teleost fish.

In conclusion, our study has revealed crucial matters regarding the molecular evolution of brain development and cognitive behavior, presenting the oHmgn2 gene as a key player in these processes within the medaka species. These insights extend our understanding of the functional diversity within protein families and underscore the intricate relationships between genetic variation, neurodevelopment, and behavior. This study emphasizes the significance of non-coding genomic regions, traditionally overlooked, as rich sources of genetic variation and functional intricacy. Further research in this direction has the potential to shed light on the detailed mechanisms of biological adaptation and the molecular mechanisms that govern the cognitive capabilities of organisms.

## Materials and methods

### Ethics statement

All procedures and experimental protocols involving live medaka fish (*Oryzias latipes*) were conducted in strict accordance with the guidelines approved by the Animal Care and Use Committee of Hokkaido University (permit no. 23-0115). To ensure the well-being of the animals, surgeries were carried out under MS-222 anesthesia, adhering to the principles outlined in the NIH Guide for the Care and Use of Laboratory Animals. Diligent efforts were taken to minimize animal discomfort and distress throughout the study.

### Fish and breeding conditions

Medaka fish (*Oryzias latipes*; d-rR strain and mutants) were kept in groups in plastic aquariums (16 cm × 25 cm × 14 cm [height]). All fish were hatched and bred within our laboratory facilities. For the experiments, we selected sexually mature medaka aged between 4 to 5 months, except for the FLAG expression assays, where one-cell stage embryos were utilized. The water temperature was maintained at approximately 28 °C, and a consistent photoperiod was provided, with 14 hours of light from 0800 to 2200 daily, using standard fluorescent lighting.

### Data reanalysis

The study included a reanalysis of data previously published by Nakayama *et al.* ^20^. In their work, total RNA and ribosome-protected fragments were extracted from specific brain regions of medaka, including the ventral telencephalon, hypothalamus, and pituitary. The Ribo-seq and RNA-seq libraries from these samples were sequenced using the Illumina HiSeq 2500 system. For our analysis, the corresponding fastq data were obtained from the NCBI Sequence Read Archive (SRA) and were processed using Trimmomatic^47^ for adapter sequence removal and quality trimming. The cleaned reads were then mapped to the medaka genome (NCBI RefSeq assembly GCF_002234675.1) using HISAT2^48^. To quantify the gene expression, the mapped reads for each exon were counted and aggregated per gene using featureCounts^49^, based on the RefSeq annotation (NCBI Annotation Release 103). From the total of 3096 lncRNA-annotated genes, we identified those with an average Transcripts Per Million (TPM) greater than 2.0 in RNA-seq and an average translation efficiency greater than 0.5 as potential peptide-coding lncRNAs, using two biological replicates for each of RNA-seq and Ribo-seq data.

### 5’- and 3’-RACE of *LOC101156433* of medaka

For the characterization of LOC101156433, total RNA was isolated from the adult medaka brain (d-rR strain) using TRIzol Reagent. The 5’ and 3’ ends of the cDNA were amplified using the SMARTer RACE cDNA Amplification Kit (Clontech) according to the manufacturer’s guidelines. The PCR-amplified products were then cloned into the pGEM-T Easy Vector (Promega) and subjected to sequencing for further analysis.

### Phylogenetic tree

To understand the evolutionary relationship of LOC101156433, its predicted amino acid sequence, along with those of medaka Hmgn1 and Hmgn3, were aligned with the corresponding sequences of Hmgn1, Hmgn2, and Hmgn3 from various species using ClustalW^50^. Phylogenetic analysis and tree construction were performed using ETE3 3.1.2^51^ on the GenomeNet platform (https://www.genome.jp/tools-bin/ete). A maximum- likelihood phylogenetic tree was inferred using PhyML^52^ v20160115 with the following model and parameters: --pinv e --alpha e --nclasses 4 -o tlr -f m --bootstrap -2. Branch supports were calculated using the Chi2-based parametric values returned by the approximate likelihood ratio test. The GenBank accession numbers of the amino acid sequences utilized in this analysis are provided in Supplementary Table 1.

The phylogenetic tree depicted in Supplementary Fig.2 was developed using the figure from Bion *et al.,* 2019^53^ as a model.

### CRISPR/Cas9 experiment for *oHmgn2* mutant generation

To create Hmgn2 mutants in medaka, we employed the CRISPR/Cas9 system. A guide RNA (gRNA) was designed using the CRISPR/Cas9 target online predictor available at COS (https://aceofbases.cos.uni-heidelberg.de:8044/index.html). The sequence “AACCTCCCCGCGGCTGGCAC<u>TGG</u>” (with the underlined section representing the PAM sequence) was selected as the target. For gRNA formation, Alt-R crRNA and Alt-R tracrRNA (IDT) were combined in equimolar amounts in IDT Duplex Buffer, heated to 95°C for 5 minutes, then gradually cooled to room temperature. Alt-R S.p. HiFi Cas9 Nuclease V3 protein (250 ng/μl) and sgRNA (50 ng/ul) were mixed and injected into fertilized eggs of the d-rR strain by a microinjector.

### *In situ* hybridization

The expression patterns of *oHmgn2* and *eomesa* were visualized using in situ hybridization on frozen brain sections, following previously described methods^30^. The cDNA fragments for *oHmgn2* and *eomesa* were amplified using synthesized cDNA, which was prepared from mRNA extracted from the medaka brain. For this, we used the ReverTra Ace qPCR RT Master Mix with gDNA Remover (TOYOBO). The primer sequences used were as follows: *oHmgn2* forward primer 5’ - CTAACCGGTTTCGTTCTTATACTTC-3’, *oHmgn2* reverse primer 5’ - CTGAAACCAGAAGTGTTTCCTCCTG -3’, *eomesa* forward primer 5’ - CTTCCAAGCTCCAGCATCAAC-3’, and *eomesa* reverse primer 5’- TATTGTTGTCCGCCTTCCCG-3’. The amplified cDNA of *oHmgn2* and *eomesa* was then cloned into the pCR2.1 TOPO vector (Thermo Fisher Scientific). From these constructs, digoxigenin (DIG) and fluorescein (FITC)-labeled riboprobes were synthesized using T7 or SP6 polymerase along with a DIG or FITC labeling mix (Roche Diagnostics). For the detection of hybridization signals, two methods were employed. Alkaline phosphatase (AP) staining was performed using anti-DIG-AP antibody (diluted at 1:1000, 11093274910; Roche Diagnostics) and 5-bromo-4-chloro-3-indolyl phosphate/nitro blue tetrazolium (BCIP/NBT) substrate (Roche Diagnostics) for chromogenic detection. For fluorescent detection, we used anti-DIG mouse monoclonal antibody [21H8] (diluted at 1:200, ab420; Abcam) and anti-FITC rabbit polyclonal antibody (diluted at 1:200, ab19491; Abcam) as the primary antibodies and Alexa Fluor 488-conjugated goat anti-rabbit secondary antibody (diluted at 1:200, A-11008; Invitrogen) and Cy3-conjugated goat anti-mouse secondary antibody (diluted at 1:200, #AP124C; Merck Millipore) for detection, with confocal microscopy (LSM900; Zeiss) utilized for imaging. The classification and naming of the brain regions in our study were consistent with those listed in the medaka brain atlases^46,54^.

### Double labeling with *in situ* hybridization and immunohistochemistry

The double labeling procedure was similar to the *in situ* hybridization protocol, with the addition of immunohistochemistry steps. A mixture of anti-FITC rabbit polyclonal antibody (diluted at 1:200, ab19491; Abcam) and mouse anti-PCNA antibody (diluted at 1:200, 610664, BD Transduction Laboratories) served as the primary antibody, and a mixture of Cy3-conjugated goat anti-rabbit antibody (diluted at 1:200, A11001; Invitrogen) and Alexa Fluor 488-conjugated goat anti-mouse antibody (diluted at 1:200, #AP124C; Merck Millipore) as the secondary antibody. This protocol was also used for immunohistochemistry with anti-PCNA antibody alone.

### Analysis of intracellular localization of Hmgn proteins

oHmgn1 CDS-FLAG, zHmgn2 CDS-FLAG and oHmgn2 CDS-FLAG were cloned into a pCS2 vector using NEBuilder HiFi DNA Assembly (New England Biolabs). The mRNAs for these constructs were synthesized from linearized plasmids using the mMessage mMachine SP6 Kit (Thermo Fisher Scientific). All synthesized RNAs were purified using the RNeasy Mini Kit (Qiagen) and subsequently microinjected into fertilized eggs of the d-rR strain medaka (concentration at 10 ng/μl). After a 24-hour incubation, the chorion was dissolved using a hatching enzyme, and the embryos were fixed with paraformaldehyde (PFA). Frozen sections of these embryos were prepared and subsequently stained for immunofluorescence. The staining utilized mouse anti-FLAG antibody (1:250 dilution, 012-22384; Fujifilm) as the primary antibody. This was followed by application of a Cy3-conjugated goat anti-mouse antibody (1:250 dilution, #AP124C; Merck Millipore) as the secondary antibody. For the visualization of these sections, confocal microscopy was employed, using an LSM900 microscope (Zeiss). The intracellular distribution of the proteins was visualized and analyzed using the “RGB Profiles Tool,” a macro in Fiji, on the fluorescent images obtained.

### Prediction of H2A-Hmgn2 complex structures by AlphaFold-Multimer

To compare the H2A-Hmgn2 complex structures in zebrafish and medaka, we utilized AlphaFold-Multimer^27^, as implemented in ColabFold v1.5.2: AlphaFold2^26^. This analysis was conducted with default parameters. The amino acid sequences used were NP_001373597 (zebrafish H2A), NP_001136015 (zebrafish Hmgn2), and XP_004083294 (medaka H2A), with accession numbers from GenBank.

### Optomotor response test

We assessed the optomotor response of *oHmgn2 ^Δ^*^2^*^/Δ^*^2^ fish as previously described^55^. Medaka were placed in a 15 cm diameter circular tank, situated within a 20 cm diameter striped tank. The water depth in the tank was maintained at 2 cm. The striped cylinder was mounted on a rotating disk, driven by a motor capable of varying speeds and directions. Medaka movements were recorded from the bottom of the tank using a video camera. Analysis of the medaka’s position and the tank’s stripes across a series of frames was performed using UMATracker software (http://ymnk13.github.io/UMATracker/).

### Shape preference test

A Y-Maze transparent plastic tank with three arms (each 13 cm long, 7 cm wide, and 10 cm high) was used for the shape preference test. One of the three arms, designated as the start zone (3 cm long, 7 cm wide, and 10 cm high), was partially blocked by an opaque plastic wall, and an opaque gate separated it from the central region. Visual cues, consisting of black vinyl tape cut into triangular and circular forms, were placed on the walls of the two unobstructed arms (the triangle zone and the circle zone), with each shape having an area of 1.5 cm². A single fish was placed in the start zone and, following a 1- minute acclimation period, the gate was opened. The movement of the fish was recorded from the bottom of the tank for 10 minutes, using a video camera, and the data from the final 9 minutes were used for quantitative analysis. Medaka positions were analyzed with UMATracker software (http://ymnk13.github.io/UMATracker/). To control for potential place preference, the triangle and circle zones were alternated between the left and right sides in different trials.

### Statistical Analysis

Statistical analyses were performed using Prism9 software (GraphPad). The Mann-Whitney U-test was employed to evaluate differences between WT fish and *oHmgn2* mutants. In the shape preference test, a preference index was calculated for each group, and its deviation from zero was assessed using a one-sample Wilcoxon signed-rank test. All statistical tests were two-tailed, with P values < 0.05 considered significant.

## Acknowledgements

We thank the National BioResource Project Medaka (https://shigen.nig.ac.jp/medaka) for providing the d-rR strain. We also thank M. Takei and M. Endo for developing the OMR apparatus. This work was supported by NIBB Collaborative Research Program (23NIBB336), KAKENHI Grant Numbers 21H05708 (SY), 23K05841 (SY), 23H03839 (SY), Astellas Foundation for Research on Metabolic Disorders (SY), Takeda Science Foundation (SY) and the Naito Foundation (SY).

