## Supplemental information for "Unique Hmgn2 Orthologous Variant Modulates Shape Preference Behavior in Medaka Fish"

**Supplementary figure legend**

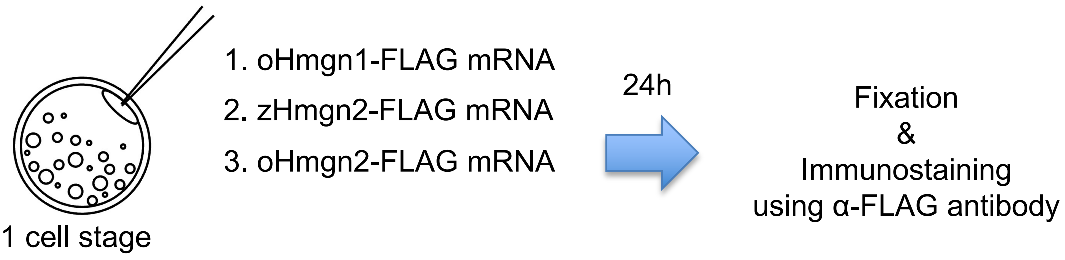


**Supplementary Fig.1 Overview of the intracellular localization analysis of oHmgn1, zHmgn2 and oHmgn2.**

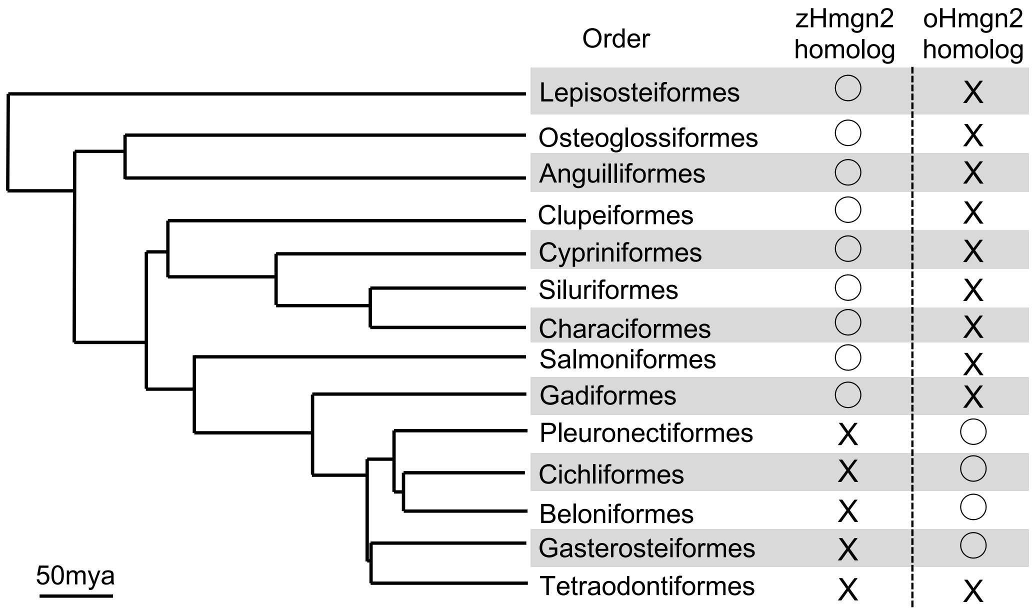


**Supplementary Fig.2 The phylogenetic tree of sequenced ray-finned fish genomes and the presence of homologs of zHmgn2 and oHmgn2.**

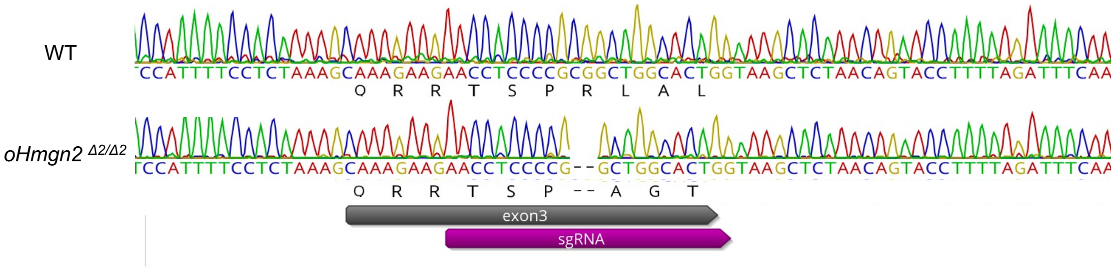


**Supplementary Fig.3 Local sequence comparison of *oHmgn2* in WT and *oHmgn2* $^{\Delta 2/\Delta 2}$ mutants.** A 2 bp deletion in exon 3 of the mutant gene lead to the incorporation of three residues with an altered sequence. In total, five residues were added to the altered sequence, followed by a stop codon in this mutant.

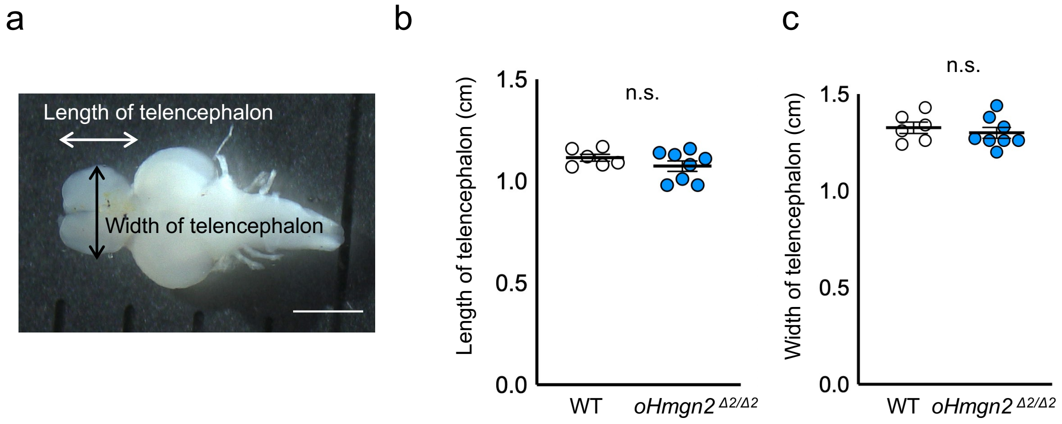


**Supplementary Fig.4 Telencephalon size comparison in WT and *oHmgn2* $^{\Delta 2/\Delta 2}$ mutants.** a. Measurements made for morphological analysis of the telencephalon. b. Telencephalon length of WT and *oHmgn2* mutants. WT, n = 6; *oHmgn2* $^{\Delta 2/\Delta 2}$, n = 8. c. Telencephalon width of WT and *oHmgn2* mutants. WT, n = 6; *oHmgn2* $^{\Delta 2/\Delta 2}$, n = 8. Mean ± SEM, Mann– Whitney U test: n.s., not significant.

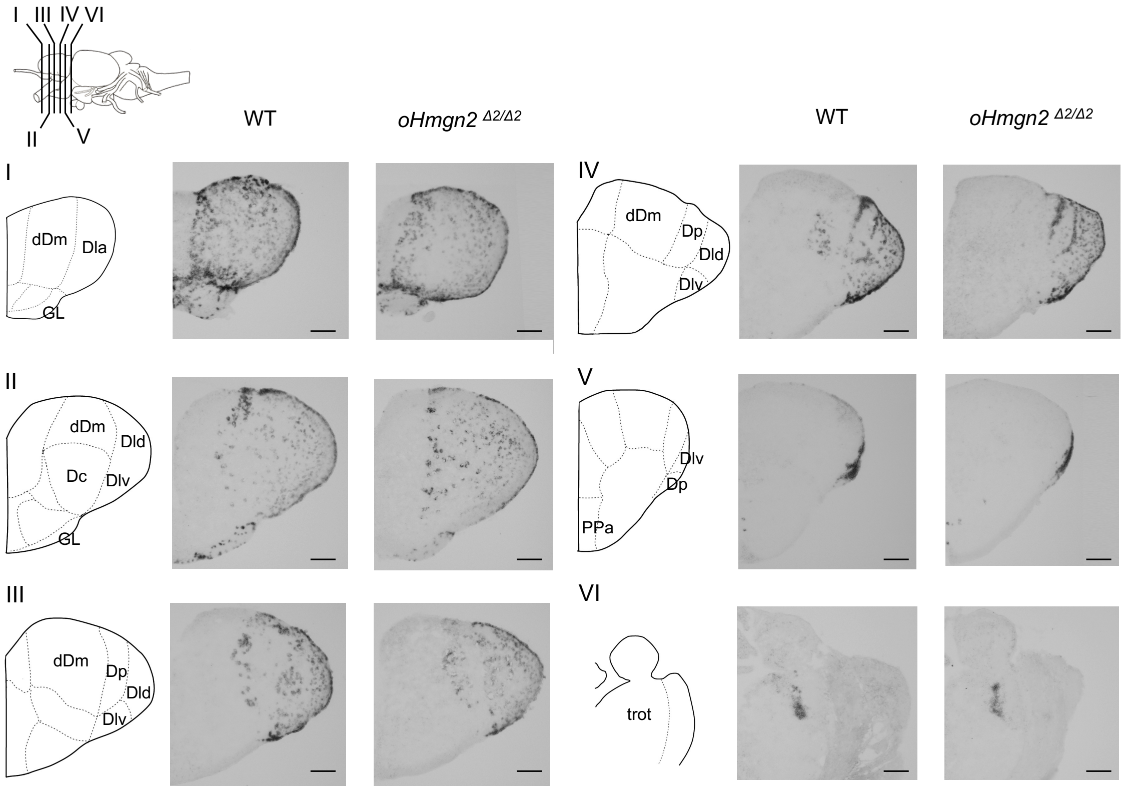


**Supplementary Fig.5 Representative micrographs showing *eomesa* expression in WT and *oHmgn2* $^{Δ2/Δ2}$ mutants.** There was no significant difference in the *eomesa* expression pattern between WT and *oHmgn2* $^{Δ2/Δ2}$ mutants. The positions of coronal sections I-VI are indicated by the lines. Micrographs II were the same as Fig. 4i. dDm, the dorsal part of dorsomedial telencephalon; Dla, the anterior part of dorsolateral telencephalon; GL, the glomerular layer of the olfactory bulb; Dc, the dorsocentral telencephalon; Dld, the dorsal part of dorsolateral telencephalon; Dlv, the ventral part of dorsolateral telencephalon; Dp, the posterior of dorsal telencephalon; PPa, the anterior parvocellular preoptic nucleus; trot, the tractus opticus dorsomedialis. Scale bars represent 100 μm.

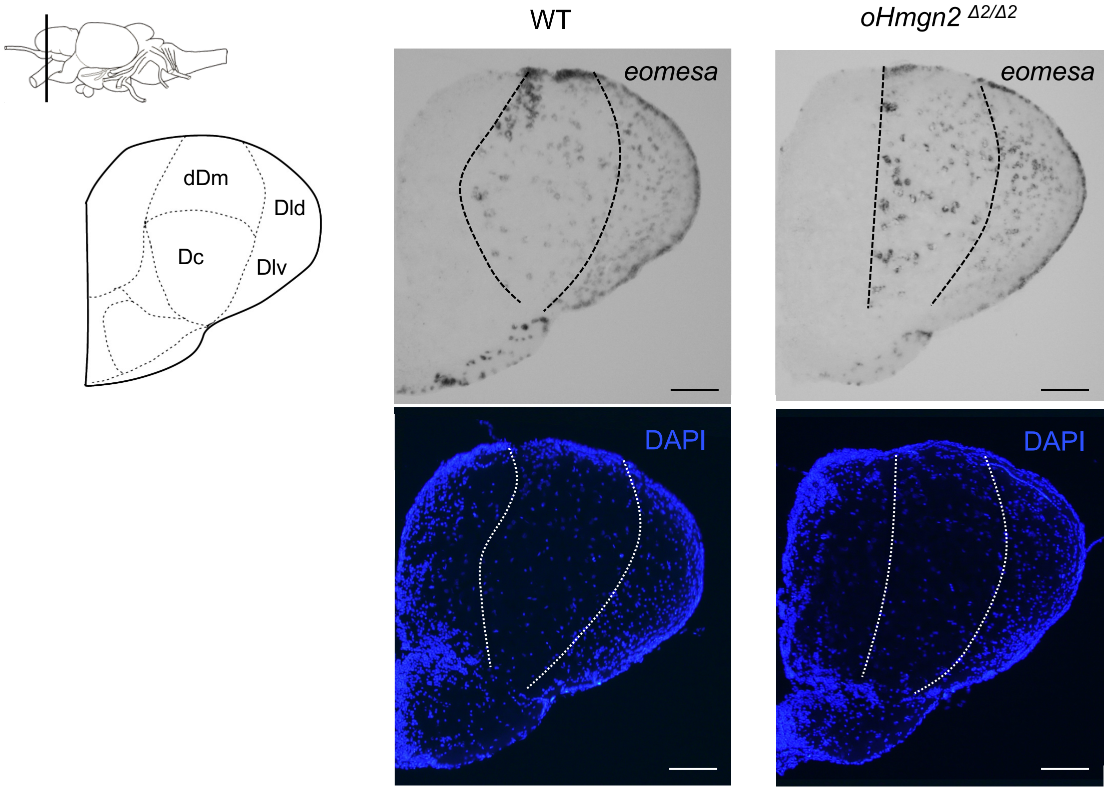


**Supplementary Fig.6 The segmentation based on DAPI staining.** Lateral view of the medaka brain (upper left). The position of the coronal section is indicated by the line. The segmentation based on *eomesa* expression identified by *in situ* hybridization (top) is almost the same as that defined by nuclear staining using DAPI (bottom) in both WT (left) and *oHmgn2* $^{\Delta 2/\Delta 2}$ mutants (right). Two serial sections were used in *in situ* hybridization (top) and nuclear staining (bottom) in both genotypes. Micrographs showing *eomesa* expression were the same as Fig. 4i. dDm, the dorsal part of dorsomedial telencephalon; Dc, the dorsocentral telencephalon; Dld, the dorsal part of dorsolateral telencephalon; Dlv, the ventral part of dorsolateral telencephalon. Scale bars represent 100 μm.

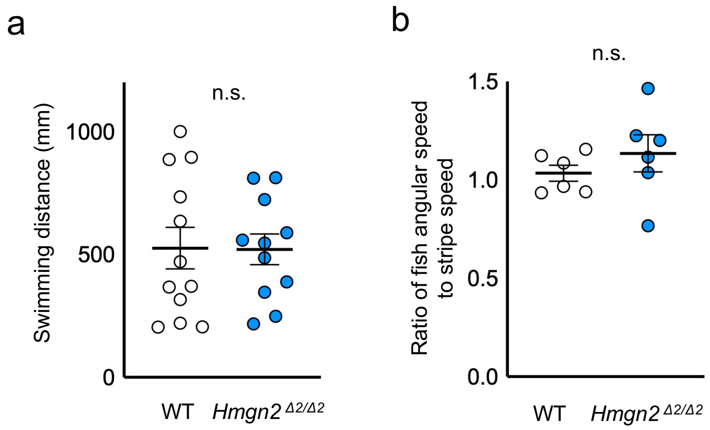


**Supplementary Fig.7 No significant defect of free swimming and the optomotor response in *oHmgn2* $^{\Delta2/\Delta2}$ mutants.** a. Two minutes after placing 1 fish in the tank, the swimming distance of it was calculated for 2 mins. Mean ± SEM, Mann– Whitney U test: n.s., not significant. WT, n = 12; *oHmgn2* $^{\Delta2/\Delta2}$, n = 11. b. Ratio of the mean fish angular speed to the mean stripe speed of WT and *oHmgn2* $^{\Delta2/\Delta2}$ mutants in the optomotor response test. Mean ± SEM, Mann– Whitney U test: n.s., not significant. n = 6 per group.

| Species | Protein | Accession NO. |
|---|---|---|
| Mouse (*Mus musculus*) | HMGN1 | NP_032277 |
| | HMGN2 | NP_058653 |
| | HMGN3 | NP_080398 |
| Chicken (*Gallus gallus*) | Hmgn1 | NP_990437 |
| | Hmgn2 | NP_001072953 |
| | Hmgn3 | NP_001006412 |
| Lizard (*Anolis carolinensis*) | Hmgn1 | XP_003218982 |
| | Hmgn2 | XP_003227420 |
| | Hmgn3 | XP_008120173 |
| Frog (*Xenopus laevis*) | Hmgn1 | NP_001080763 |
| | Hmgn2 | NP_001079778 |
| | Hmgn3 | NP_001079722 |
| Zebrafish (*Danio rerio*) | Hmgn1 | NP_001154808 |
| | Hmgn2 | NP_001136015 |
| | Hmgn3 | NP_001230102 |
| Medaka (*Oryzias latipes*) | Hmgn1 | XP_004073635 |
| | Hmgn3 | XP_004083393 |

**Supplementary Table 1. Species names and GenBank accession numbers of Hmgns used to construct the phylogenetic tree.**

**Supplementary Dataset 1. The list of lncRNA-annotated genes predicted to be expressed in the adult medaka brain.** Yellow highlighted genes are identified as potentially translated (translation efficiency > 0.5).

**Supplementary Movie 1. Predicted zH2A-zHmgn2 complex structures in zebrafish.** zH2A is shown in light blue, zHmgn2 in green. The nucleosome binding core domain in zHmgn2, along with the essential positively charged residues in zH2A necessary for binding, are represented as spherical models. The atomic structures are color-coded, with oxygen atoms depicted in red and nitrogen atoms in blue.

**Supplementary Movie 2. Predicted oH2A-oHmgn2 complex structures in medaka.** oH2A is shown in light blue, oHmgn2 in green. The altered nucleosome binding core domain in oHmgn2, along with the essential positively charged residues in oH2A necessary for binding, are represented as spherical models. The atomic structures are color-coded, with oxygen atoms depicted in red and nitrogen atoms in blue.
