## Supplemental dataset for "Unique Hmgn2 Orthologous Variant Modulates Shape Preference Behavior in Medaka Fish"

| gene | TPM | | | | | | Translation efficiency |
|---|---|---|---|---|---|---|---|
| | RNA-seq_1 | RNA-seq_2 | Ribo-seq_1 | Ribo-seq_2 | RNA-seq_average | Ribo-seq_average | |
| LOC105354757 | 303.470395 | 430.281858 | 15.2818479 | 9.93417127 | 366.8761262 | 12.60800961 | 0.034365849 |
| LOC111946391 | 387.978649 | 342.974119 | 611.855443 | 1006.22072 | 365.4763838 | 809.0380799 | 2.21365351 |
| LOC101159365 | 321.178104 | 389.336448 | 407.452025 | 522.026875 | 355.2572759 | 464.7394501 | 1.308177148 |
| LOC101156433 | 269.529035 | 385.651565 | 281.251369 | 312.592368 | 327.5902998 | 296.9218687 | 0.906381748 |
| LOC101155702 | 269.782198 | 307.78053 | 392.024596 | 774.938566 | 288.7813638 | 583.481581 | 2.020495967 |
| LOC111948018 | 244.357264 | 269.137488 | 193.083208 | 215.720743 | 256.747376 | 204.4019753 | 0.796120991 |
| LOC110016053 | 240.458072 | 228.770027 | 184.117844 | 124.794024 | 234.6140497 | 154.4559339 | 0.658340513 |
| LOC101166873 | 193.742512 | 242.400791 | 14.1106678 | 19.4540648 | 218.0716515 | 16.78236628 | 0.076958037 |
| LOC110015861 | 172.793255 | 230.828332 | 46.7013287 | 49.3838011 | 201.8107938 | 48.04256488 | 0.23805746 |
| LOC111947724 | 140.410768 | 207.498079 | 6.70893529 | 3.96583044 | 173.9544236 | 5.337382865 | 0.030682651 |
| LOC101170841 | 160.079751 | 165.926695 | 93.1430407 | 117.713058 | 163.0032227 | 105.4280494 | 0.646785061 |
| LOC110016374 | 141.802068 | 167.778585 | 48.1512006 | 74.190534 | 154.7903265 | 61.17086728 | 0.395185337 |
| LOC110015771 | 137.728431 | 152.782269 | 118.664421 | 212.755952 | 145.2553499 | 165.7101864 | 1.140819849 |
| LOC110016932 | 121.437199 | 162.926473 | 6.28628292 | 2.90280867 | 142.1818357 | 4.594545791 | 0.032314576 |
| LOC111947627 | 92.8501603 | 120.691562 | 59.4086076 | 43.2278131 | 106.7708611 | 51.31821035 | 0.480638723 |
| LOC101161693 | 83.7476207 | 120.699285 | 3.35412521 | 1.27167097 | 102.2234528 | 2.31289809 | 0.022625905 |
| LOC105354512 | 113.800983 | 82.2382556 | 36.866211 | 26.5898562 | 98.01961946 | 31.72803361 | 0.323690643 |
| LOC105354516 | 70.3027171 | 112.529399 | 9.30109229 | 8.30929251 | 91.4160578 | 8.805192404 | 0.096319975 |
| LOC110016232 | 76.1900195 | 101.689454 | 74.3439469 | 323.763812 | 88.93973693 | 199.0538796 | 2.238075876 |
| LOC111946921 | 86.3046696 | 85.109717 | 31.9729583 | 46.0640271 | 85.70719333 | 39.01849269 | 0.455253418 |
| LOC101168817 | 62.0444379 | 95.3195356 | 63.6856057 | 131.061848 | 78.68198675 | 97.37372667 | 1.237560599 |
| LOC110016708 | 72.7196754 | 76.4936716 | 49.2620394 | 58.7548312 | 74.60667347 | 54.0084353 | 0.723908905 |
| LOC110014943 | 72.7688068 | 76.2022982 | 55.4670809 | 82.7468631 | 74.48555253 | 69.10697199 | 0.927790285 |
| LOC110016179 | 68.1856304 | 71.9111014 | 82.8859094 | 104.226467 | 70.04836588 | 93.55618842 | 1.33559416 |
| LOC101163943 | 64.5650058 | 74.8129488 | 81.7201011 | 97.0647406 | 69.68897733 | 89.39242083 | 1.282734003 |
| LOC110015770 | 65.5772875 | 71.1585336 | 20.5652931 | 29.9053568 | 68.36791056 | 25.23532496 | 0.369110665 |
| LOC101172299 | 62.3882994 | 73.4703187 | 79.4238232 | 103.627765 | 67.92930905 | 91.52579411 | 1.347368248 |
| LOC105354749 | 62.4291618 | 73.1987368 | 20.2989598 | 24.3987872 | 67.81394929 | 22.34887352 | 0.329561598 |
| LOC110015180 | 66.4043758 | 66.6350152 | 31.692006 | 40.4606708 | 66.5196955 | 36.07633839 | 0.542340703 |
| LOC105353949 | 61.1487222 | 66.423699 | 16.4326134 | 12.0450571 | 63.7862106 | 14.23883528 | 0.223227484 |
| LOC111948625 | 58.1898264 | 67.7746895 | 51.2586132 | 86.8778783 | 62.98225797 | 69.06824574 | 1.096630193 |
| LOC110016898 | 54.8473406 | 68.5559053 | 91.5914017 | 103.324741 | 61.70162295 | 97.45807129 | 1.5795058 |
| LOC105356364 | 52.8468019 | 61.8959024 | 28.1635112 | 35.2749157 | 57.37135216 | 31.71921342 | 0.552875472 |
| LOC101173718 | 52.5677501 | 60.703895 | 69.9786278 | 88.1510139 | 56.63582254 | 79.06482084 | 1.39602141 |
| LOC110017602 | 51.5321364 | 61.2646883 | 29.4000044 | 30.3998507 | 56.39841238 | 29.89992752 | 0.530155482 |
| LOC101162358 | 49.0383367 | 63.0651358 | 7.45804946 | 7.97533295 | 56.05173624 | 7.716691201 | 0.137670868 |
| LOC101175482 | 46.3724364 | 59.0636074 | 11.032976 | 14.7426426 | 52.71802192 | 12.88780931 | 0.24446686 |
| LOC110016370 | 53.1716482 | 50.6434233 | 34.0873509 | 30.215262 | 51.90753573 | 32.15130642 | 0.619395738 |
| LOC105354508 | 46.4416131 | 57.1085974 | 35.220963 | 57.7550201 | 51.77510526 | 46.48799155 | 0.897883091 |
| LOC101156488 | 43.6525132 | 57.3631945 | 4.8689225 | 4.30730012 | 50.50785386 | 4.588111308 | 0.090839562 |
| LOC110015154 | 42.9187884 | 57.9799932 | 2.98896739 | 3.18098623 | 50.4493908 | 3.084976808 | 0.061149932 |
| LOC110017634 | 48.3097769 | 51.3292545 | 11.0059028 | 16.0300234 | 49.81951567 | 13.51796313 | 0.27133871 |
| LOC101163334 | 44.1387417 | 54.7194829 | 15.7022751 | 18.4689962 | 49.42911231 | 17.08563562 | 0.345659366 |
| LOC101163932 | 37.5977088 | 57.1638549 | 21.6503562 | 31.6718276 | 47.38078189 | 26.6610919 | 0.562698437 |
| LOC105356609 | 37.7416318 | 53.9167618 | 0.52083713 | 0.5485231 | 45.8291968 | 0.534680119 | 0.011666801 |
| LOC111947337 | 38.7040016 | 51.1923963 | 7.49426764 | 12.8537247 | 44.94819895 | 10.17399619 | 0.226349363 |
| LOC111949072 | 42.1115759 | 47.1725287 | 15.1778376 | 13.3335298 | 44.64205233 | 14.25568369 | 0.319333072 |
| LOC105357036 | 41.9086292 | 47.3743049 | 1.38121864 | 0.4134239 | 44.64146706 | 0.897321268 | 0.020100622 |
| LOC111948498 | 87.8428962 | 0.97834445 | 51.0178008 | 1.47541613 | 44.41062032 | 26.24660845 | 0.590998465 |
| LOC110016062 | 42.1825035 | 46.6369068 | 1.18259812 | 0.74727666 | 44.40970514 | 0.964937393 | 0.021728075 |
| LOC105354195 | 37.8950531 | 49.4236983 | 15.914468 | 28.5597696 | 43.65937569 | 22.23711879 | 0.509332038 |
| LOC105354127 | 44.7193716 | 42.1883036 | 15.1029516 | 15.9057743 | 43.45383762 | 15.50436294 | 0.356800775 |
| LOC105354678 | 35.7728528 | 45.595375 | 18.124019 | 15.9939513 | 40.68411387 | 17.05898513 | 0.419303347 |
| LOC101172898 | 35.3543462 | 43.8018839 | 1.96213553 | 2.83396955 | 39.57811504 | 2.398052539 | 0.060590368 |
| LOC101156819 | 31.4979418 | 44.2433552 | 18.5015664 | 27.3619826 | 37.87064852 | 22.93177451 | 0.605528963 |
| LOC111947342 | 31.5058265 | 41.1564383 | 0.76596464 | 0.26889359 | 36.33113239 | 0.517429114 | 0.014242031 |
| LOC111947352 | 29.861946 | 41.9481425 | 0.93512308 | 0 | 35.90504424 | 0.467561541 | 0.013022169 |
| LOC101161305 | 29.4935711 | 40.0062496 | 18.4331055 | 26.3692545 | 34.74991036 | 22.40118002 | 0.644639937 |
| LOC105356807 | 29.8791777 | 38.686581 | 3.98085847 | 4.31225246 | 34.28287935 | 4.146555462 | 0.120951202 |
| LOC111946424 | 34.5744602 | 33.7034752 | 10.9882011 | 22.7588515 | 34.13896768 | 16.87352631 | 0.494260004 |
| LOC111947381 | 31.1894498 | 36.6997692 | 14.1570306 | 9.26147483 | 33.94460947 | 11.70925269 | 0.344951757 |
| LOC111947072 | 27.6158282 | 40.1335294 | 5.12613389 | 2.92160727 | 33.8746788 | 4.023870582 | 0.118786974 |
| LOC105358699 | 32.517241 | 34.9642258 | 12.137955 | 2.50445746 | 33.74073341 | 7.32120621 | 0.216984205 |
| LOC110014995 | 28.2149882 | 38.7479566 | 0.55747722 | 0.70453303 | 33.4814724 | 0.631005125 | 0.018846397 |
| LOC105356303 | 28.6150626 | 38.0691089 | 3.6774845 | 2.98771766 | 33.34208573 | 3.33260108 | 0.099951788 |
| LOC110017301 | 23.1936083 | 42.2567738 | 9.70441034 | 11.7533042 | 32.72519106 | 10.72885728 | 0.327847048 |
| LOC111946746 | 30.3940427 | 34.4762326 | 3.33041184 | 3.78804109 | 32.43513763 | 3.559226464 | 0.109733663 |
| LOC110015921 | 29.9606678 | 33.9336949 | 9.11091891 | 12.5377608 | 31.94718136 | 10.82433984 | 0.33881987 |
| LOC101167910 | 35.061272 | 28.6851269 | 30.6540738 | 42.6799374 | 31.87319945 | 36.66700557 | 1.150402413 |
| LOC105356658 | 29.9515791 | 33.2897623 | 6.68844999 | 6.54408999 | 31.6206707 | 6.616269989 | 0.209238762 |
| LOC110017659 | 27.7577853 | 33.7232465 | 1.05840016 | 0.51679746 | 30.7405159 | 0.78759881 | 0.025620872 |
| LOC110016392 | 31.1138331 | 29.5600786 | 7.2569974 | 11.584808 | 30.33695583 | 9.420902673 | 0.310542123 |
| LOC110017601 | 26.3120766 | 34.0844732 | 1.21305129 | 22.2290754 | 30.19827493 | 11.72106332 | 0.38813685 |
| LOC110016890 | 30.4610953 | 29.0979349 | 12.5000912 | 41.2489374 | 29.77951512 | 26.87451431 | 0.902449694 |
| LOC111946715 | 26.9733123 | 31.7929114 | 26.5568679 | 42.55214 | 29.38311185 | 34.55450392 | 1.175998788 |
| LOC110016696 | 27.8371089 | 30.1430248 | 5.43540291 | 15.0263685 | 28.99006682 | 10.2308857 | 0.352910042 |
| LOC101165207 | 23.7039903 | 33.5464024 | 38.248424 | 42.3741328 | 28.62519636 | 40.31127838 | 1.408244606 |
| LOC110016936 | 23.8878996 | 32.3889756 | 24.4593131 | 33.1337739 | 28.13843764 | 28.79654348 | 1.023388144 |
| LOC105356393 | 27.7153874 | 27.5067679 | 39.0602516 | 15.3303968 | 27.61107763 | 27.19532421 | 0.984942514 |
| LOC110015834 | 30.1411694 | 24.0432858 | 1.87593552 | 5.92696203 | 27.09222762 | 3.901448775 | 0.144006201 |
| LOC111946320 | 23.584867 | 29.6486154 | 5.4677565 | 5.58122258 | 26.6167412 | 5.524489539 | 0.207556947 |
| LOC111948911 | 25.9075704 | 26.9519272 | 12.973683 | 21.0805522 | 26.42974877 | 17.02711759 | 0.644240615 |
| LOC111948956 | 19.9069349 | 32.8097332 | 10.1404125 | 9.0872352 | 26.35833405 | 9.613823836 | 0.36473564 |
| LOC110017121 | 24.0880197 | 28.4734111 | 1.29800666 | 2.25555723 | 26.28071544 | 1.776781948 | 0.067607823 |
| LOC111948010 | 25.74578 | 26.7684828 | 0.44510386 | 0.18750564 | 26.2571314 | 0.316304751 | 0.012046432 |
| LOC111947900 | 24.4302569 | 27.6036196 | 6.53977617 | 4.13244535 | 26.01693825 | 5.33611076 | 0.205101412 |
| LOC111949080 | 25.0596312 | 26.1699264 | 12.0155781 | 1.88573678 | 25.6147788 | 6.950657434 | 0.271353405 |
| LOC101171961 | 22.313295 | 28.7560621 | 18.0649343 | 31.6294052 | 25.53467855 | 24.84716975 | 0.973075486 |
| LOC110013812 | 22.384557 | 28.568978 | 7.00969587 | 6.88136566 | 25.47676754 | 6.945530765 | 0.272622135 |
| LOC101170603 | 28.3181826 | 22.6032359 | 5.01517531 | 4.52722744 | 25.46070925 | 4.771201375 | 0.187394677 |
| LOC101166127 | 22.5994942 | 28.1933818 | 12.9682319 | 24.1634107 | 25.39643802 | 18.5658213 | 0.731040364 |
| LOC110016042 | 21.5003959 | 28.5421054 | 14.582386 | 16.2609216 | 25.02125066 | 15.42165379 | 0.616342244 |
| LOC110016634 | 23.680209 | 24.6890047 | 4.78511768 | 8.56711367 | 24.18460686 | 6.676115676 | 0.276048137 |
| LOC110016246 | 22.8268514 | 25.4571569 | 12.2142034 | 12.8634698 | 24.14200411 | 12.53883657 | 0.519378446 |
| LOC110013348 | 20.5738957 | 27.565472 | 0.84940932 | 0.17891222 | 24.06968385 | 0.51416077 | 0.021361343 |
| LOC105354499 | 25.2292751 | 22.0860299 | 12.1430927 | 12.9649733 | 23.65765253 | 12.55403303 | 0.530654215 |
| LOC110015643 | 21.9065835 | 25.3905104 | 10.3880288 | 8.75217752 | 23.64854695 | 9.570103177 | 0.404680389 |
| LOC110015646 | 23.960309 | 23.3199393 | 24.4939482 | 32.1690858 | 23.64012415 | 28.331517 | 1.198450432 |
| LOC110017482 | 19.5194918 | 27.4988998 | 7.43791977 | 13.8388209 | 23.5091958 | 10.63837033 | 0.452519534 |
| LOC110016707 | 23.4447654 | 23.2212932 | 16.0852574 | 15.0042623 | 23.33302929 | 15.54475986 | 0.666212675 |
| LOC105355641 | 21.7332184 | 24.879957 | 6.05659181 | 14.6706422 | 23.30658769 | 10.36361701 | 0.444664708 |
| LOC111948927 | 23.5333218 | 22.6172013 | 0.58822258 | 0.55754149 | 23.07526151 | 0.572882035 | 0.024826676 |
| LOC111946611 | 24.5985432 | 21.2861854 | 4.90696096 | 7.44163011 | 22.94236432 | 6.174295534 | 0.269122025 |
| LOC110014811 | 20.3798776 | 25.1246779 | 9.26691643 | 7.31963617 | 22.75227771 | 8.293276302 | 0.364503124 |
| LOC110013545 | 22.6464364 | 22.057165 | 14.1693353 | 10.8384692 | 22.35180071 | 12.50390223 | 0.559413642 |
| LOC110016173 | 24.0004918 | 20.694624 | 2.92892177 | 1.54230676 | 22.34755787 | 2.235614266 | 0.100038415 |
| LOC111948623 | 23.415921 | 20.836964 | 86.4326123 | 101.657983 | 22.12644246 | 94.0452975 | 4.250357809 |
| LOC105356231 | 18.1181702 | 24.9152932 | 3.4668677 | 5.53438211 | 21.51673172 | 4.500624906 | 0.209168612 |
| LOC110017592 | 19.8381774 | 22.7324238 | 8.4296424 | 8.77329023 | 21.28530061 | 8.601466318 | 0.404103587 |
| LOC110015155 | 17.8635451 | 24.6697038 | 0.94782345 | 0.11978479 | 21.26662445 | 0.533804118 | 0.025100557 |
| LOC105353883 | 20.6511384 | 21.7272296 | 0.92467051 | 0.58429375 | 21.18918399 | 0.754482127 | 0.035606946 |
| LOC105355754 | 22.3636103 | 19.3218685 | 46.9290636 | 59.1398982 | 20.84273938 | 53.03448087 | 2.544506262 |
| LOC101159004 | 19.2397297 | 22.3349843 | 24.8119919 | 43.1728159 | 20.78735696 | 33.9924039 | 1.635244152 |
| LOC110017753 | 19.5779175 | 21.8796185 | 2.09363668 | 1.85213905 | 20.72876797 | 1.972887862 | 0.09517632 |
| LOC110014720 | 17.7639565 | 23.5875596 | 0.59440789 | 0.46950348 | 20.67575806 | 0.531955683 | 0.025728473 |
| LOC110016061 | 25.0523115 | 16.1118356 | 0.26790612 | 0.33857654 | 20.58207356 | 0.303241326 | 0.014733274 |
| LOC110015606 | 18.2715141 | 22.8584605 | 46.002759 | 61.8585503 | 20.56498728 | 53.93065463 | 2.622450182 |
| LOC105356023 | 19.1901416 | 21.3318966 | 43.0267065 | 62.1893708 | 20.26101908 | 52.60803863 | 2.596514935 |
| LOC105354424 | 16.5137074 | 23.7970277 | 11.6793782 | 21.1563699 | 20.15536753 | 16.41787403 | 0.814565847 |
| LOC111949208 | 19.6915552 | 20.6064585 | 0 | 0 | 20.14900686 | 0 | 0 |
| LOC105354109 | 19.7021232 | 20.0774873 | 25.8000458 | 27.3851988 | 19.88980524 | 26.5926223 | 1.336997622 |
| LOC110015327 | 20.3348787 | 19.3638086 | 3.95139488 | 5.4343483 | 19.84934368 | 4.69287159 | 0.236424522 |
| LOC111948347 | 16.1042302 | 22.9057865 | 6.91241457 | 14.3170492 | 19.50500835 | 10.6147319 | 0.544205453 |
| LOC110015232 | 18.812386 | 20.1699428 | 5.66540882 | 7.95541749 | 19.49116442 | 6.810413155 | 0.349410277 |
| LOC101173743 | 18.0957457 | 20.8459853 | 4.74664193 | 1.6360225 | 19.47086552 | 3.191332213 | 0.163902946 |
| LOC111946843 | 15.2079724 | 23.3292992 | 2.62921815 | 5.81485516 | 19.26863577 | 4.222036653 | 0.219114456 |
| LOC110015479 | 17.0508633 | 21.2358294 | 1.40094647 | 0.22131242 | 19.14334635 | 0.811129447 | 0.042371351 |
| LOC101166651 | 16.39128 | 21.7210825 | 1.19737747 | 1.13492347 | 19.05618124 | 1.166150466 | 0.061195391 |
| LOC101161157 | 19.3363854 | 18.4699623 | 3.25810895 | 2.57347439 | 18.90317382 | 2.915791668 | 0.154248789 |
| LOC110016034 | 14.1266716 | 23.2757916 | 1.56788238 | 0.45726253 | 18.70123159 | 1.012572456 | 0.054144694 |
| LOC111946402 | 14.3500617 | 22.9528389 | 1.17993975 | 1.77796172 | 18.6514503 | 1.478950734 | 0.079294141 |
| LOC105355140 | 15.7318806 | 21.5587987 | 12.4022588 | 11.1366658 | 18.64533965 | 11.76946234 | 0.631228101 |
| LOC111947755 | 18.4351569 | 18.4808287 | 0.47216998 | 0.23868911 | 18.45799282 | 0.355429544 | 0.019256132 |
| LOC105355110 | 20.1138906 | 16.7303018 | 5.00741843 | 3.51573076 | 18.42209619 | 4.261574593 | 0.231329516 |
| LOC110014080 | 15.1800863 | 21.525356 | 0.79393526 | 0.41806911 | 18.35272117 | 0.606002182 | 0.033019745 |
| LOC101159265 | 16.939016 | 19.4266946 | 6.27533195 | 8.882372 | 18.18285531 | 7.578851974 | 0.416813083 |
| LOC110014794 | 18.1547335 | 18.0903924 | 10.9450938 | 15.8664369 | 18.12256295 | 13.40576532 | 0.739727894 |
| LOC111947859 | 14.4135946 | 21.8078231 | 19.6619792 | 19.4129481 | 18.11070886 | 19.53746366 | 1.078779622 |
| LOC105353661 | 15.0483894 | 21.1351387 | 1.24878182 | 0 | 18.09176407 | 0.624390909 | 0.034512439 |
| LOC110015156 | 17.3518873 | 18.6238871 | 3.20777877 | 4.72961107 | 17.98788721 | 3.968694915 | 0.220631521 |
| LOC110017270 | 17.072874 | 18.870975 | 1.78886678 | 0.45214968 | 17.97192449 | 1.120508229 | 0.062347704 |
| LOC111946408 | 16.7870074 | 18.985469 | 0.40972353 | 0.25890185 | 17.88623819 | 0.33431269 | 0.018691057 |
| LOC105354200 | 14.8928915 | 20.8606428 | 1.92354539 | 2.279019 | 17.87676716 | 2.101282199 | 0.117542628 |
| LOC111948236 | 20.1441527 | 15.179678 | 12.0974726 | 11.8139497 | 17.66191538 | 11.95571118 | 0.67692042 |
| LOC110015089 | 14.8903435 | 20.3040453 | 2.485536 | 5.60926893 | 17.59719441 | 4.047402465 | 0.230002714 |
| LOC110015765 | 17.5319789 | 17.1825118 | 6.06200431 | 11.0128145 | 17.35724534 | 8.537409422 | 0.491864305 |
| LOC110015781 | 17.01233 | 17.3981439 | 10.703905 | 16.0867173 | 17.20523697 | 13.39531115 | 0.778560108 |
| LOC110016506 | 16.6543552 | 17.7300856 | 5.94735468 | 11.8111644 | 17.19222036 | 8.879259552 | 0.516469622 |
| LOC101156352 | 15.5658313 | 18.7993929 | 13.5323198 | 21.57182 | 17.18261214 | 17.55206989 | 1.021501838 |
| LOC105354181 | 13.6494241 | 20.6952057 | 1.89793816 | 3.54991541 | 17.17231489 | 2.723926787 | 0.158623156 |
| LOC105355439 | 13.5951369 | 20.6230496 | 0.64439361 | 1.34685413 | 17.10909324 | 0.995623872 | 0.058192673 |
| LOC105353806 | 16.2225248 | 17.9033385 | 2.0984194 | 1.73397206 | 17.06293165 | 1.916195731 | 0.112301671 |
| LOC111947157 | 16.4251538 | 17.622076 | 6.89878061 | 8.45439634 | 17.02361492 | 7.676588475 | 0.450937625 |
| LOC110014636 | 17.0867164 | 16.6587538 | 2.36321866 | 0.44799111 | 16.87273511 | 1.405604883 | 0.083306285 |
| LOC111949008 | 15.5040921 | 17.7111558 | 15.7402951 | 12.3246368 | 16.60762394 | 14.03246595 | 0.844941215 |
| LOC110014902 | 14.6170161 | 18.589811 | 5.44287939 | 7.86130939 | 16.60341355 | 6.652094388 | 0.40064619 |
| LOC105353621 | 14.8699987 | 17.6699367 | 9.61703634 | 8.36566814 | 16.26996774 | 8.99135224 | 0.552634915 |
| LOC105354005 | 13.7837991 | 18.5015181 | 21.9576529 | 28.7178354 | 16.14265861 | 25.33774419 | 1.569614077 |
| LOC110014949 | 15.7370726 | 16.3238794 | 7.92887157 | 10.0204128 | 16.03047598 | 8.974642194 | 0.559848766 |
| LOC110016535 | 17.4210539 | 14.1972342 | 17.2238742 | 13.3815526 | 15.80914405 | 15.3027134 | 0.967965967 |
| LOC101168047 | 14.7783686 | 16.1525741 | 10.0601017 | 14.2879298 | 15.46547135 | 12.17401573 | 0.787173921 |
| LOC105354433 | 14.1638036 | 16.476707 | 6.61148424 | 8.00736807 | 15.32025531 | 7.309426154 | 0.477108639 |
| LOC110017176 | 14.6923426 | 15.9361816 | 1.50095276 | 3.86672937 | 15.31426208 | 2.683841067 | 0.175251086 |
| LOC110013716 | 13.3399792 | 16.8354375 | 0 | 0.74416301 | 15.08770838 | 0.372081506 | 0.024661234 |
| LOC101167112 | 14.3268007 | 15.5617549 | 2.39995582 | 6.182726 | 14.94427779 | 4.291340909 | 0.287156126 |
| LOC110015460 | 12.1530173 | 17.7249202 | 2.14881615 | 1.90095366 | 14.93896872 | 2.024884907 | 0.135543821 |
| LOC110015116 | 11.3250022 | 18.496365 | 9.81879235 | 14.4031551 | 14.91068359 | 12.1109737 | 0.812234639 |
| LOC110016714 | 14.0064789 | 15.6159555 | 0.693692 | 0.73056636 | 14.81121724 | 0.712129179 | 0.048080395 |
| LOC110017506 | 13.3276388 | 16.1539431 | 2.04812284 | 16.1774568 | 14.74079094 | 9.112789798 | 0.618202228 |
| LOC111947081 | 11.0165568 | 18.4017795 | 0.76183545 | 5.77679103 | 14.70916814 | 3.269313238 | 0.222263639 |
| LOC110017271 | 15.6734081 | 13.4806799 | 3.5017463 | 0.42147286 | 14.577044 | 1.961609577 | 0.134568406 |
| LOC111947106 | 9.3543756 | 19.7462757 | 0.99521462 | 1.41495784 | 14.55032565 | 1.205086228 | 0.082821942 |
| LOC110015530 | 14.8395815 | 14.0790518 | 11.8694363 | 18.7505641 | 14.45931664 | 15.31000016 | 1.058832899 |
| LOC111948195 | 13.320649 | 15.5956898 | 1.14352374 | 0.12043097 | 14.45816939 | 0.631977353 | 0.043710745 |
| LOC110014081 | 15.1606406 | 13.699147 | 1.29256533 | 2.58642022 | 14.42989383 | 1.939492773 | 0.134407972 |
| LOC105356551 | 13.3404711 | 15.3332976 | 3.67384252 | 3.40483596 | 14.33688435 | 3.53933924 | 0.246869484 |
| LOC110013904 | 15.9169249 | 12.6912414 | 2.23874021 | 1.41464652 | 14.30408314 | 1.826693362 | 0.127704331 |
| LOC110015987 | 13.2877954 | 15.1511128 | 5.06979285 | 8.40937573 | 14.21945408 | 6.739584287 | 0.473969271 |
| LOC110016907 | 8.98273092 | 19.3936075 | 0 | 0 | 14.1881692 | 0 | 0 |
| LOC111949061 | 11.5673846 | 16.6492291 | 7.11046418 | 12.3559141 | 14.10830685 | 9.73318916 | 0.689890663 |
| LOC110015540 | 13.4566453 | 14.6165931 | 2.60099035 | 3.72538047 | 14.0366192 | 3.16318541 | 0.22535237 |
| LOC105354754 | 11.8230648 | 16.2301028 | 4.43395351 | 5.78241485 | 14.0265838 | 5.108184179 | 0.36417878 |
| LOC110015442 | 13.0561499 | 14.6584413 | 5.88835315 | 5.39518183 | 13.85729559 | 5.641767489 | 0.407133373 |
| LOC105354276 | 10.8299861 | 16.8541644 | 14.6530447 | 20.5759358 | 13.84207523 | 17.61449024 | 1.272532474 |
| LOC110014937 | 13.4800217 | 14.1533476 | 13.4235808 | 9.42475581 | 13.81668461 | 11.4241683 | 0.82683861 |
| LOC110014724 | 13.9018455 | 13.670306 | 0.22185318 | 0 | 13.78607575 | 0.11092659 | 0.008046277 |
| LOC111947057 | 14.5105653 | 12.9159856 | 1.67243166 | 1.40906606 | 13.71327545 | 1.54074886 | 0.112354548 |
| LOC110015471 | 12.9621066 | 13.8122662 | 4.88933116 | 4.51548015 | 13.38718641 | 4.702405652 | 0.351261685 |
| LOC110017661 | 12.5546657 | 14.1097144 | 0.55221287 | 0.11631333 | 13.33219005 | 0.334263099 | 0.025071882 |
| LOC110014471 | 11.6794462 | 14.7268016 | 1.65206596 | 4.43670495 | 13.20312391 | 3.044385452 | 0.230580692 |
| LOC110017218 | 12.1458229 | 14.0674644 | 6.07177124 | 8.24893907 | 13.10664364 | 7.160355154 | 0.546314934 |
| LOC101157765 | 11.8246511 | 14.1757894 | 2.11276459 | 1.40921235 | 13.00022028 | 1.760988466 | 0.135458356 |
| LOC105357648 | 5.43037904 | 20.5292422 | 0.09547391 | 0.27751519 | 12.97981059 | 0.186494547 | 0.014368048 |
| LOC110017568 | 11.8648043 | 14.0598927 | 2.65684057 | 2.37012936 | 12.9623485 | 2.513484963 | 0.193906603 |
| LOC111947746 | 10.6613552 | 15.1929923 | 6.16407462 | 7.10272329 | 12.9271217 | 6.633398955 | 0.513138122 |
| LOC105355259 | 10.6881343 | 15.0112544 | 5.32169155 | 8.03322413 | 12.84969435 | 6.677457842 | 0.519658885 |
| LOC110016558 | 10.6051609 | 14.9255457 | 0.41382277 | 0.34865617 | 12.76535329 | 0.381239469 | 0.029865172 |
| LOC111947988 | 12.59482 | 12.8804514 | 0.85390016 | 0.43165951 | 12.73763572 | 0.642779833 | 0.050463041 |
| LOC110015402 | 13.4678649 | 11.8885289 | 0.29411129 | 0.92923581 | 12.67819687 | 0.611673551 | 0.0482461 |
| LOC110017605 | 10.8335778 | 14.4668125 | 0.64677563 | 0.20434682 | 12.65019513 | 0.425561227 | 0.033640685 |
| LOC101162141 | 10.5304801 | 14.695601 | 2.50326124 | 2.44459347 | 12.61304052 | 2.473927354 | 0.196140443 |
| LOC110014889 | 10.676124 | 14.2829837 | 0.41790185 | 0.26406967 | 12.47955384 | 0.340985762 | 0.027323554 |
| LOC105354588 | 12.786851 | 12.1257947 | 3.3762571 | 5.18120322 | 12.45632288 | 4.278730156 | 0.343498655 |
| LOC105353870 | 10.2257224 | 14.4797902 | 0.9791257 | 0 | 12.35275628 | 0.489562849 | 0.039631871 |
| LOC110015913 | 11.1214764 | 13.3640912 | 6.34500094 | 8.01873612 | 12.24278378 | 7.181868531 | 0.586620548 |
| LOC110016523 | 12.1908491 | 12.2485942 | 2.66223816 | 4.76638161 | 12.21972166 | 3.714309884 | 0.303960269 |
| LOC110013528 | 11.4485786 | 12.8790365 | 0.71254021 | 0.90049978 | 12.16380759 | 0.806519996 | 0.066304896 |
| LOC111947878 | 11.1322949 | 13.078919 | 8.24827679 | 9.81600236 | 12.10560698 | 9.032139573 | 0.746112077 |
| LOC110014500 | 9.48563219 | 14.5227632 | 4.14294554 | 7.63554881 | 12.00419767 | 5.889247174 | 0.490598983 |
| LOC105354971 | 12.0353581 | 11.8647594 | 0 | 0.21321025 | 11.95005878 | 0.106605127 | 0.008920887 |
| LOC110017029 | 4.49136546 | 19.3936075 | 0 | 0 | 11.94248647 | 0 | 0 |
| LOC110016499 | 11.3752188 | 12.4448095 | 1.64645214 | 4.50832737 | 11.91001417 | 3.077389755 | 0.258386742 |
| LOC111948213 | 11.6903968 | 11.9555173 | 11.193701 | 7.66266863 | 11.82295703 | 9.428184834 | 0.797447272 |
| LOC110014175 | 10.3512656 | 13.1503312 | 0.6669206 | 0.58999238 | 11.75079839 | 0.628456492 | 0.053482025 |
| LOC105356473 | 13.5777822 | 9.64938871 | 15.7553233 | 19.609701 | 11.61358545 | 17.68251213 | 1.52257132 |
| LOC111947692 | 10.6574774 | 12.5340487 | 0 | 0.69856241 | 11.59576304 | 0.349281205 | 0.030121451 |
| LOC105355961 | 10.4333883 | 12.6355283 | 3.07589911 | 3.02344343 | 11.53445829 | 3.049671268 | 0.264396575 |
| LOC110017701 | 10.5794526 | 12.4665079 | 1.04827427 | 0.86939768 | 11.52298026 | 0.958835975 | 0.083210763 |
| LOC105356660 | 10.3835514 | 12.5018422 | 0.45833668 | 0 | 11.44269678 | 0.229168339 | 0.020027476 |
| LOC101163508 | 10.2366738 | 12.4702193 | 4.06003576 | 2.96020645 | 11.35344658 | 3.510121108 | 0.309167889 |
| LOC105354595 | 11.3346531 | 11.3644175 | 8.29940803 | 3.78758335 | 11.34953533 | 6.04349569 | 0.532488381 |
| LOC111947543 | 10.4798527 | 12.1465226 | 4.34832233 | 0 | 11.31318766 | 2.174161163 | 0.19217936 |
| LOC110016937 | 10.4057378 | 12.2182759 | 12.3930262 | 11.8729261 | 11.31200685 | 12.13297613 | 1.072575034 |
| LOC110014733 | 10.2913662 | 12.2788852 | 3.05008221 | 8.03053609 | 11.28512571 | 5.540309148 | 0.490939072 |
| LOC105356634 | 12.2420981 | 10.1768568 | 4.15866875 | 3.66020272 | 11.20947744 | 3.909435737 | 0.34876164 |
| LOC105356423 | 9.98354865 | 12.4325368 | 0.56359113 | 0 | 11.20804273 | 0.281795565 | 0.025142264 |
| LOC111948211 | 9.92798816 | 12.4505657 | 1.65375025 | 1.61499207 | 11.18927693 | 1.634371156 | 0.146065842 |
| LOC105354397 | 13.1250564 | 9.12744471 | 3.26752545 | 2.40885137 | 11.12625057 | 2.838188406 | 0.255089384 |
| LOC105354219 | 8.69604802 | 13.5251681 | 7.14119425 | 10.2124498 | 11.11060806 | 8.676822038 | 0.780949341 |
| LOC105355332 | 11.1407444 | 11.0071826 | 0.7834815 | 0.49507726 | 11.07396349 | 0.639279382 | 0.057728146 |
| LOC110015800 | 11.2427413 | 10.8856643 | 0.35474233 | 0 | 11.06420278 | 0.177371165 | 0.016031084 |
| LOC110016489 | 7.60651326 | 14.0300136 | 3.38154677 | 1.06838957 | 10.81826344 | 2.224968171 | 0.205667775 |
| LOC111949030 | 10.3786525 | 11.2119978 | 4.85220517 | 7.6651984 | 10.79532511 | 6.258701783 | 0.57976038 |
| LOC110015328 | 8.18313177 | 13.3447042 | 0.91766543 | 0 | 10.76391798 | 0.458832713 | 0.042626924 |
| LOC110014262 | 9.23239826 | 12.2332564 | 5.16315232 | 6.52513 | 10.73282733 | 5.844141157 | 0.544510871 |
| LOC110017172 | 8.67998884 | 12.7363325 | 28.9687402 | 20.7788453 | 10.70816069 | 24.87379272 | 2.322881907 |
| LOC111949132 | 9.96864041 | 11.2403258 | 1.32570803 | 1.34033113 | 10.60448309 | 1.333019577 | 0.1257034 |
| LOC105355928 | 9.30139541 | 11.7505339 | 2.27020844 | 0.71726555 | 10.52596465 | 1.493736998 | 0.141909749 |
| LOC110017752 | 11.0943067 | 9.87094058 | 13.3494912 | 13.0895122 | 10.48262361 | 13.21950171 | 1.261087129 |
| LOC101175589 | 11.4031878 | 9.51356945 | 8.0932431 | 6.29424221 | 10.45837862 | 7.193742651 | 0.687844924 |
| LOC110016026 | 9.2650321 | 11.6285919 | 1.74739383 | 2.48437729 | 10.44681202 | 2.115885555 | 0.202538875 |
| LOC105353928 | 9.77559891 | 11.0973986 | 0.53755699 | 0.20586609 | 10.43649875 | 0.37171154 | 0.035616498 |
| LOC110017389 | 8.23813065 | 12.5321265 | 0 | 0.26999112 | 10.38512856 | 0.134995558 | 0.01299893 |
| LOC111946663 | 8.79585919 | 11.7938849 | 5.67714967 | 5.89351481 | 10.29487205 | 5.785332236 | 0.56196252 |
| LOC111947461 | 9.98839333 | 10.4909937 | 6.53397179 | 13.4480185 | 10.23969354 | 9.990995135 | 0.97571232 |
| LOC110015529 | 7.13159654 | 13.3291899 | 0 | 0 | 10.23039324 | 0 | 0 |
| LOC110013478 | 9.19066451 | 11.2402546 | 0.56079554 | 0 | 10.21635957 | 0.280397769 | 0.027445957 |
| LOC110016690 | 8.27614539 | 12.1532002 | 0.14675023 | 0 | 10.21467278 | 0.073375117 | 0.007183306 |
| LOC105356432 | 9.92700855 | 10.473699 | 0.09318858 | 1.05993545 | 10.20035377 | 0.576562011 | 0.056523727 |
| LOC110014805 | 11.3169017 | 8.9027322 | 9.03006234 | 5.70604226 | 10.10981694 | 7.368052297 | 0.728801752 |
| LOC110015951 | 4.59010038 | 15.3809856 | 2.63393001 | 19.7163189 | 9.985542979 | 11.17512446 | 1.119130375 |
| LOC110017394 | 9.6760004 | 10.2912549 | 2.40887174 | 9.13290968 | 9.983627661 | 5.770890711 | 0.57803545 |
| LOC110013481 | 9.60307982 | 10.1421063 | 0 | 0.17648135 | 9.872593059 | 0.088240673 | 0.008937943 |
| LOC105355733 | 8.22699704 | 11.4867941 | 0.45991115 | 0.25832453 | 9.856895563 | 0.35911784 | 0.036433159 |
| LOC105353864 | 9.31595905 | 10.1666424 | 1.81595838 | 5.57353848 | 9.741300698 | 3.694748429 | 0.37928697 |
| LOC101167053 | 7.20115596 | 12.2741121 | 1.48050022 | 1.44580242 | 9.737634028 | 1.463151321 | 0.150257374 |
| LOC105355476 | 8.18519874 | 11.2541114 | 1.13207324 | 1.19225049 | 9.719655054 | 1.162161861 | 0.119568221 |
| LOC110017171 | 9.54266767 | 9.45219158 | 2.49022865 | 3.85522335 | 9.497429626 | 3.172725998 | 0.334061543 |
| LOC105353802 | 9.39875897 | 9.52207766 | 0 | 0.11515095 | 9.460418316 | 0.057575475 | 0.006085933 |
| LOC111946619 | 9.17936782 | 9.68461608 | 0.60264595 | 0.15232335 | 9.43199195 | 0.377484649 | 0.040021731 |
| LOC110017170 | 8.53622091 | 10.2817756 | 2.59736881 | 3.03384753 | 9.408998248 | 2.815608169 | 0.299246327 |
| LOC110016054 | 7.84576777 | 10.9090671 | 0.8018183 | 0.17733246 | 9.37741743 | 0.489575382 | 0.052207912 |
| LOC110015331 | 8.96303195 | 9.77738659 | 0 | 3.91664743 | 9.370209273 | 1.958323713 | 0.208994661 |
| LOC110017427 | 9.01358267 | 9.52022648 | 0.30655201 | 0.96854188 | 9.266904577 | 0.637546944 | 0.068798264 |
| LOC110014653 | 6.81190428 | 11.6106466 | 0 | 1.48832602 | 9.211275431 | 0.744163011 | 0.08078827 |
| LOC110017497 | 8.28374897 | 10.1356785 | 0.30688486 | 4.2662114 | 9.209713708 | 2.286548127 | 0.248275701 |
| LOC110016099 | 6.98486279 | 11.3611991 | 0.27601655 | 0.0872066 | 9.173030945 | 0.181611579 | 0.019798427 |
| LOC105358491 | 8.51195729 | 9.7572478 | 0.38810979 | 0.49048849 | 9.134602544 | 0.439299139 | 0.048091763 |
| LOC111948034 | 7.90485215 | 10.3233861 | 0.84692195 | 0.09730271 | 9.114119135 | 0.472112328 | 0.051800105 |
| LOC101155654 | 8.03018188 | 10.1809949 | 0.57053079 | 0 | 9.105588388 | 0.285265393 | 0.031328606 |
| LOC105354098 | 8.16241254 | 10.0170284 | 0.92366324 | 0.38910484 | 9.08972048 | 0.656384041 | 0.072211686 |
| LOC110016399 | 7.40104195 | 10.7696592 | 3.97225121 | 5.50278378 | 9.08535058 | 4.737517492 | 0.521445755 |
| LOC110016885 | 8.36722347 | 9.66435322 | 0 | 0.25809122 | 9.015788346 | 0.129045609 | 0.014313292 |
| LOC111949281 | 8.62887978 | 9.39852545 | 0.29108234 | 0 | 9.013702616 | 0.14554117 | 0.016146658 |
| LOC105358620 | 8.26421929 | 9.75667179 | 0.37819931 | 0.71694563 | 9.010445541 | 0.54757247 | 0.060770854 |
| LOC105356723 | 8.73321062 | 9.16425616 | 0.78773955 | 0.69687506 | 8.948733391 | 0.742307306 | 0.082951103 |
| LOC111948287 | 4.06485145 | 13.8120878 | 0 | 0.34379042 | 8.9384696 | 0.17189521 | 0.019230944 |
| LOC105354197 | 3.74967208 | 14.0605995 | 7.13084969 | 14.3370855 | 8.905135806 | 10.73396761 | 1.20536821 |
| LOC105353893 | 7.5024728 | 10.2882826 | 1.0612301 | 0.22352831 | 8.895377691 | 0.642379209 | 0.072214945 |
| LOC110017769 | 9.008092 | 8.75556955 | 1.23558886 | 1.85430783 | 8.881830778 | 1.544948344 | 0.173944807 |
| LOC111949078 | 8.72854793 | 9.00088566 | 7.54514018 | 6.00733835 | 8.864716795 | 6.776239264 | 0.764405612 |
| LOC110013436 | 8.723158 | 8.98780987 | 2.10116774 | 1.79884008 | 8.855483937 | 1.950003913 | 0.220202976 |
| LOC105354472 | 8.01038043 | 9.6414053 | 0.63916995 | 0 | 8.825892867 | 0.319584974 | 0.036209931 |
| LOC110017283 | 8.27062428 | 9.29471374 | 8.29633212 | 3.69351087 | 8.78266901 | 5.994921498 | 0.682585384 |
| LOC110015262 | 7.25818455 | 10.2314121 | 3.4592485 | 6.04331084 | 8.74479833 | 4.751279673 | 0.543326386 |
| LOC105356558 | 8.62613005 | 8.8217576 | 4.82904138 | 8.07734725 | 8.723943825 | 6.453194316 | 0.739710668 |
| LOC110016174 | 8.45391462 | 8.97816108 | 0 | 0 | 8.716037851 | 0 | 0 |
| LOC110017643 | 6.96177585 | 10.4322772 | 13.1390932 | 1.01404754 | 8.697026523 | 7.076570364 | 0.81367699 |
| LOC111947940 | 8.34110729 | 8.84620692 | 4.4863643 | 3.64488005 | 8.593657102 | 4.065622179 | 0.473095695 |
| LOC111947186 | 6.68968881 | 10.4594493 | 0.1334471 | 0 | 8.574569072 | 0.066723549 | 0.007781563 |
| LOC111948977 | 7.48128216 | 9.66435322 | 0.81688136 | 3.61327705 | 8.572817692 | 2.215079205 | 0.258384032 |
| LOC110014662 | 8.78955391 | 8.13280314 | 4.77580716 | 3.29214973 | 8.461178524 | 4.033978444 | 0.476763188 |
| LOC110017179 | 8.95775441 | 7.93033368 | 2.49633814 | 3.85591825 | 8.444044043 | 3.176128194 | 0.376138279 |
| LOC110016620 | 8.30241693 | 8.55755909 | 0.67812128 | 0 | 8.42998801 | 0.339060642 | 0.040220774 |
| LOC105357133 | 6.58310137 | 10.262799 | 1.1658731 | 2.84158887 | 8.422950175 | 2.003730987 | 0.23788945 |
| LOC110017584 | 8.44160255 | 8.39323849 | 3.40531267 | 4.91839236 | 8.417420523 | 4.161852513 | 0.494433241 |
| LOC101162274 | 7.90616309 | 8.90572723 | 2.85255796 | 5.70796187 | 8.40594516 | 4.280259913 | 0.509194366 |
| LOC105355285 | 7.62778793 | 9.18319035 | 0.80125004 | 0 | 8.405489139 | 0.40062502 | 0.047662309 |
| LOC101158341 | 8.11781339 | 8.66286584 | 2.92892177 | 1.38807608 | 8.390339619 | 2.158498928 | 0.257260019 |
| LOC110015238 | 7.71158975 | 9.04316021 | 0 | 0.13479179 | 8.37737498 | 0.067395896 | 0.00804499 |
| LOC105356142 | 6.57890152 | 10.0921516 | 0 | 0 | 8.33552654 | 0 | 0 |
| LOC111947658 | 5.96179629 | 10.6132977 | 0.91617812 | 0 | 8.287546987 | 0.458089062 | 0.055274385 |
| LOC110016224 | 8.74566884 | 7.75147304 | 1.61278717 | 1.27388818 | 8.248570937 | 1.443337673 | 0.174980331 |
| LOC105355772 | 6.30355322 | 10.1896421 | 0.63277825 | 1.19954635 | 8.246597644 | 0.916162298 | 0.111095792 |
| LOC111947433 | 7.38322529 | 9.10873951 | 0.243132 | 0 | 8.245982399 | 0.121566001 | 0.014742452 |
| LOC110016368 | 8.46950799 | 7.91916214 | 0 | 0 | 8.194335067 | 0 | 0 |
| LOC110017364 | 4.1919411 | 12.0572099 | 0 | 0 | 8.124575504 | 0 | 0 |
| LOC110014883 | 8.45149415 | 7.7788451 | 0 | 0 | 8.115169626 | 0 | 0 |
| LOC110013479 | 7.46919329 | 8.75890882 | 0.41321777 | 0 | 8.114051058 | 0.206608883 | 0.025463099 |
| LOC111947751 | 7.27222036 | 8.86127712 | 4.49400717 | 4.46244259 | 8.066748738 | 4.47822488 | 0.555146196 |
| LOC110015620 | 7.26215808 | 8.61514928 | 1.50661488 | 0.38080836 | 7.93865368 | 0.943711621 | 0.118875525 |
| LOC105355429 | 7.81693934 | 7.99421568 | 0 | 0 | 7.90557751 | 0 | 0 |
| LOC101160133 | 5.69483959 | 9.85598295 | 0 | 1.72282112 | 7.775411272 | 0.861410559 | 0.110786495 |
| LOC105355528 | 5.10423682 | 10.4399864 | 0.31145008 | 0 | 7.772111584 | 0.155725042 | 0.020036388 |
| LOC110014269 | 6.62583628 | 8.91103588 | 0.18830177 | 0.11898676 | 7.768436081 | 0.153644262 | 0.019778017 |
| LOC110014929 | 7.67538511 | 7.84945121 | 7.96171693 | 31.4435075 | 7.762418159 | 19.70261222 | 2.538205469 |
| LOC110015717 | 7.60688625 | 7.90693003 | 3.36323329 | 4.74084628 | 7.756908138 | 4.052039781 | 0.522378209 |
| LOC105354487 | 8.33770414 | 7.10565886 | 0.8072161 | 2.55037507 | 7.721681501 | 1.678795584 | 0.217413213 |
| LOC105355867 | 7.69383001 | 7.70090347 | 0.25470798 | 0.10729896 | 7.69736674 | 0.181003469 | 0.023514986 |
| LOC111947148 | 6.20270959 | 9.17524266 | 0.45957878 | 1.74243046 | 7.688976127 | 1.101004623 | 0.143192618 |
| LOC110015319 | 6.64912887 | 8.67645711 | 0.16373118 | 0.07759556 | 7.662792992 | 0.120663373 | 0.015746657 |
| LOC110017776 | 7.46217997 | 7.84945121 | 0.88463522 | 2.79497845 | 7.655815588 | 1.83980683 | 0.240314936 |
| LOC111948735 | 7.5196346 | 7.69016852 | 0 | 0 | 7.604901556 | 0 | 0 |
| LOC110015780 | 5.74305748 | 9.46254101 | 0.6619226 | 0.20913246 | 7.602799241 | 0.435527532 | 0.057285155 |
| LOC105354748 | 6.84280728 | 8.35244181 | 1.83176248 | 2.19921149 | 7.597624544 | 2.015486987 | 0.265278572 |
| LOC105357936 | 6.23040026 | 8.92037481 | 0.86171022 | 1.3612738 | 7.575387535 | 1.111492009 | 0.146724112 |
| LOC105356859 | 6.56569088 | 8.5797549 | 0.22702084 | 0.14345311 | 7.572722889 | 0.185236978 | 0.02446108 |
| LOC110016681 | 6.09293764 | 8.97280738 | 9.10113975 | 13.4188965 | 7.532872509 | 11.26001813 | 1.49478411 |
| LOC110017282 | 5.63312146 | 9.38808789 | 5.62684895 | 8.75217752 | 7.510604677 | 7.189513238 | 0.957248257 |
| LOC111947071 | 7.74841368 | 7.23896631 | 0.54388894 | 0.53461358 | 7.493689995 | 0.539251259 | 0.071960711 |
| LOC105355112 | 6.75131555 | 8.14368096 | 0.35913717 | 0.11346831 | 7.447498255 | 0.23630274 | 0.031729143 |
| LOC110015954 | 6.36957284 | 8.50442165 | 0.73413234 | 3.01530986 | 7.436997244 | 1.874721102 | 0.252080381 |
| LOC111949043 | 8.80694749 | 6.06572038 | 7.45754489 | 7.77542354 | 7.436333933 | 7.616484215 | 1.024225685 |
| LOC111946459 | 4.11457306 | 10.7534848 | 0 | 0 | 7.434028903 | 0 | 0 |
| LOC110016091 | 7.29463766 | 7.56977588 | 0.44510386 | 0 | 7.432206769 | 0.22255193 | 0.02994426 |
| LOC105358615 | 6.31567285 | 8.53497861 | 0 | 2.36555129 | 7.425325729 | 1.182775647 | 0.159289396 |
| LOC105354057 | 8.25685368 | 6.55529329 | 2.02852357 | 2.42120023 | 7.406073481 | 2.2248619 | 0.300410454 |
| LOC111947979 | 7.13050114 | 7.68009362 | 0.9442348 | 1.5910835 | 7.40529738 | 1.267659151 | 0.171182747 |
| LOC110013435 | 6.89353829 | 7.85380621 | 0.87815653 | 0.71344523 | 7.373672249 | 0.795800879 | 0.107924634 |
| LOC105357891 | 6.14407053 | 8.59503068 | 0.89524283 | 0.32325633 | 7.369550606 | 0.609249582 | 0.082671199 |
| LOC110015859 | 6.95744262 | 7.76866608 | 0 | 0.09306885 | 7.363054352 | 0.046534425 | 0.006319989 |
| LOC105355117 | 5.65537923 | 9.02247007 | 0 | 0 | 7.33892465 | 0 | 0 |
| LOC105354014 | 7.60184711 | 7.02725393 | 1.01023374 | 1.48950817 | 7.314550521 | 1.249870956 | 0.170874608 |
| LOC110016884 | 6.10178299 | 8.49953124 | 5.45638902 | 6.3440615 | 7.300657115 | 5.900225259 | 0.808177287 |
| LOC111946844 | 7.17042556 | 7.33303995 | 0.76286357 | 2.89229348 | 7.251732753 | 1.827578525 | 0.252019564 |
| LOC111946708 | 9.04706038 | 5.44249059 | 0 | 0 | 7.24477548 | 0 | 0 |
| LOC111947533 | 7.99974524 | 6.44576821 | 1.00583968 | 1.9067522 | 7.222756726 | 1.456295942 | 0.201626055 |
| LOC110013912 | 5.93244529 | 8.4441066 | 0.35986879 | 0 | 7.188275946 | 0.179934397 | 0.025031649 |
| LOC105356044 | 4.70150819 | 9.67028649 | 0.87674587 | 1.03876962 | 7.18589734 | 0.957757745 | 0.133282971 |
| LOC101160165 | 7.41431759 | 6.90545744 | 2.19740292 | 4.51270864 | 7.159887514 | 3.355055781 | 0.468590571 |
| LOC111947170 | 6.58331689 | 7.66770217 | 1.13815148 | 3.35622512 | 7.125509531 | 2.247188302 | 0.315372296 |
| LOC111948158 | 5.77687996 | 8.36951202 | 0.99873128 | 0 | 7.07319599 | 0.499365638 | 0.070599717 |
| LOC110015423 | 5.98848728 | 8.11469365 | 0 | 0 | 7.051590468 | 0 | 0 |
| LOC110017134 | 6.59976886 | 7.47259607 | 6.8459746 | 3.24444773 | 7.036182462 | 5.045211163 | 0.717038137 |
| LOC110016093 | 7.47648031 | 6.58408617 | 3.87769598 | 6.26185948 | 7.030283241 | 5.069777729 | 0.721134207 |
| LOC105355382 | 6.41120403 | 7.54009049 | 1.33007506 | 0.84046646 | 6.975647258 | 1.085270762 | 0.155579937 |
| LOC110017382 | 5.19002231 | 8.71983253 | 3.07636409 | 1.70094403 | 6.954927422 | 2.38865406 | 0.343447734 |
| LOC111947930 | 6.29671824 | 7.51277132 | 0.19792784 | 0 | 6.904744781 | 0.098963919 | 0.014332741 |
| LOC110015128 | 5.95826783 | 7.85110164 | 2.6148448 | 7.36026662 | 6.904684737 | 4.987555707 | 0.722343727 |
| LOC101175339 | 7.13760062 | 6.66165277 | 10.7823354 | 10.810425 | 6.899626695 | 10.7963802 | 1.56477744 |
| LOC105356939 | 5.20044928 | 8.53741165 | 1.56155222 | 3.40872258 | 6.868930463 | 2.485137399 | 0.361793938 |
| LOC110016928 | 7.19957363 | 6.51328955 | 2.01065717 | 2.54104443 | 6.856431589 | 2.2758508 | 0.331929338 |
| LOC105356836 | 6.67677899 | 6.99077437 | 0 | 0 | 6.833776682 | 0 | 0 |
| LOC110014078 | 5.9212133 | 7.69766624 | 0.72872437 | 0.39469419 | 6.80943977 | 0.561709281 | 0.082489794 |

| | | | | | | | |
|---|---|---|---|---|---|---|---|
| LOC110016529 | 6.78162915 | 6.68773243 | 1.25618201 | 2.38132164 | 6.734680792 | 1.81875182 | 0.270057613 |
| LOC110015992 | 6.97538999 | 6.46480802 | 2.71335313 | 8.00124069 | 6.720099001 | 5.357296912 | 0.797205058 |
| LOC105355958 | 6.20005659 | 7.17496244 | 0.33849216 | 1.06945582 | 6.687509515 | 0.703973991 | 0.105266989 |
| LOC111948785 | 5.78617352 | 7.53123022 | 0 | 1.03435785 | 6.658701869 | 0.517178927 | 0.077669632 |
| LOC110015184 | 6.7063415 | 6.5409112 | 2.31884099 | 1.30245486 | 6.623626351 | 1.810647922 | 0.273362026 |
| LOC110017192 | 6.17393137 | 7.01549642 | 2.27706708 | 2.33815568 | 6.594713899 | 2.307611382 | 0.349918346 |
| LOC111946716 | 6.17106311 | 6.97852451 | 0.49240584 | 0.38893537 | 6.574793813 | 0.440670605 | 0.067024247 |
| LOC105355199 | 6.33907187 | 6.80166024 | 0 | 0 | 6.570366056 | 0 | 0 |
| LOC110017712 | 7.05896817 | 6.06400609 | 2.34313742 | 5.92245795 | 6.561487129 | 4.132797686 | 0.629856861 |
| LOC111947069 | 6.18188448 | 6.927064 | 0.36818187 | 0.1938766 | 6.554474241 | 0.281029236 | 0.042875939 |
| LOC110015440 | 6.02413304 | 7.01377834 | 0.38454551 | 0 | 6.518955691 | 0.192272756 | 0.029494411 |
| LOC105353894 | 5.23992637 | 7.7609627 | 0.37485537 | 0 | 6.500444536 | 0.187427687 | 0.028833057 |
| LOC105356873 | 6.17509737 | 6.76622059 | 0.34314968 | 1.1564492 | 6.470658978 | 0.749799437 | 0.115876828 |
| LOC105356114 | 4.7322187 | 8.15080126 | 0 | 0 | 6.441509977 | 0 | 0 |
| LOC110017063 | 6.87145691 | 5.99662177 | 0 | 0.60053505 | 6.43403934 | 0.300267523 | 0.046668587 |
| LOC110016729 | 5.82889562 | 7.0219574 | 0 | 0 | 6.425426506 | 0 | 0 |
| LOC110016999 | 5.38490457 | 7.43448516 | 2.23431582 | 0.70592539 | 6.409694867 | 1.470120603 | 0.229358906 |
| LOC105355414 | 6.03027332 | 6.75623754 | 0.39842254 | 0.30211298 | 6.393255426 | 0.350267761 | 0.054787074 |
| LOC110013818 | 5.30412367 | 7.43485999 | 0.3847549 | 0.14587459 | 6.369491833 | 0.265314746 | 0.04165399 |
| LOC105357319 | 5.44507846 | 7.27322722 | 0.46107822 | 0 | 6.359152843 | 0.230539112 | 0.036253117 |
| LOC111948944 | 7.51238326 | 5.19104914 | 2.10611737 | 1.86318012 | 6.351716202 | 1.984648744 | 0.312458662 |
| LOC110016274 | 5.4328071 | 7.26555798 | 1.30049517 | 3.56102545 | 6.34918254 | 2.430760313 | 0.382846185 |
| LOC110013444 | 7.51905335 | 5.12638272 | 1.11422188 | 3.52035064 | 6.322718035 | 2.317286264 | 0.366501598 |
| LOC105354846 | 5.75722024 | 6.87732901 | 1.10406622 | 0.82449879 | 6.317274626 | 0.964282503 | 0.152642169 |
| LOC105354772 | 5.57470989 | 7.00139492 | 0.6087027 | 1.28211861 | 6.288052409 | 0.945410654 | 0.150350314 |
| LOC105355048 | 4.88636936 | 7.65193924 | 1.26716409 | 2.08185483 | 6.269154299 | 1.67450946 | 0.267102927 |
| LOC105356738 | 4.04190292 | 8.49467012 | 1.00009046 | 0.48611628 | 6.26828652 | 0.743103374 | 0.118549682 |
| LOC110015342 | 6.6689972 | 5.84591996 | 4.54597334 | 6.11983008 | 6.257458579 | 5.332901708 | 0.852247225 |
| LOC105354736 | 4.21354904 | 8.29901886 | 3.88509899 | 4.90994152 | 6.256283952 | 4.39752025 | 0.702896525 |
| LOC110015138 | 6.56104095 | 5.92044302 | 3.20273033 | 7.08325132 | 6.240741984 | 5.142990824 | 0.824099256 |
| LOC111949293 | 4.90948056 | 7.53123022 | 0 | 0 | 6.22035539 | 0 | 0 |
| LOC105356454 | 5.92339503 | 6.5120583 | 0.92165528 | 4.46497807 | 6.217726664 | 2.693316671 | 0.433167429 |
| LOC110017494 | 7.74597811 | 4.65979127 | 0.94528746 | 1.79196444 | 6.202884692 | 1.368625952 | 0.220643462 |
| LOC110017732 | 5.18297065 | 7.19355277 | 5.37632244 | 5.58122258 | 6.188261712 | 5.478772511 | 0.885349193 |
| LOC105356286 | 5.7226499 | 6.65048969 | 0.2698243 | 0.34100071 | 6.186569793 | 0.305412504 | 0.049367018 |
| LOC110017642 | 4.99385418 | 7.37511969 | 0.49650009 | 0.25098834 | 6.184486935 | 0.373744217 | 0.060432534 |
| LOC111948205 | 6.11182277 | 6.25042964 | 0.69161734 | 0 | 6.181126205 | 0.345808668 | 0.0559459 |
| LOC110013510 | 4.70906375 | 7.63894745 | 1.34751347 | 1.06435711 | 6.1740056 | 1.205935291 | 0.195324619 |
| LOC110017195 | 4.60908129 | 7.73681909 | 0.46161611 | 0.33336281 | 6.172950193 | 0.397489459 | 0.064392138 |
| LOC110016083 | 5.60550263 | 6.72960553 | 0 | 0 | 6.167554081 | 0 | 0 |
| LOC101156115 | 5.52914309 | 6.78544281 | 7.34132341 | 16.2362839 | 6.157292947 | 11.78880364 | 1.914608212 |
| LOC110016667 | 6.40283431 | 5.82048079 | 6.64169337 | 3.73052998 | 6.111657552 | 5.186111673 | 0.848560579 |
| LOC111948424 | 5.963047 | 6.17002434 | 0.72770585 | 0 | 6.066535673 | 0.363852924 | 0.059977052 |
| LOC111946660 | 5.21218855 | 6.90604388 | 0.06800793 | 0 | 6.059116214 | 0.034003965 | 0.005612034 |
| LOC110014974 | 5.49663746 | 6.55817501 | 0.13575454 | 0 | 6.027406238 | 0.06787727 | 0.011261439 |
| LOC105354203 | 4.06890266 | 7.96363607 | 2.61974105 | 1.01163252 | 6.016269365 | 1.815686784 | 0.301796125 |
| LOC110017606 | 5.60189497 | 6.41640995 | 0 | 0.88124567 | 6.009152461 | 0.440622836 | 0.073325288 |
| LOC111948469 | 3.53865158 | 8.4441066 | 0 | 4.63893825 | 5.99137909 | 2.319469125 | 0.387134429 |
| LOC111947780 | 6.28229242 | 5.37887327 | 1.9701514 | 3.06443535 | 5.830582845 | 2.517293375 | 0.431739578 |
| LOC110017615 | 4.95611468 | 6.701699 | 0.57122262 | 0.07219043 | 5.828906837 | 0.321706524 | 0.055191571 |
| LOC110017504 | 5.83093519 | 5.82286203 | 5.40803431 | 8.27347396 | 5.826898609 | 6.840754135 | 1.173995738 |
| LOC111948042 | 5.72772733 | 5.8576235 | 0.84877163 | 2.94983536 | 5.792675413 | 1.899303492 | 0.327880186 |
| LOC110014830 | 5.91571726 | 5.50809676 | 1.09905749 | 2.54645541 | 5.711907011 | 1.82275645 | 0.319115218 |
| LOC105357134 | 5.13555911 | 6.28408329 | 0.18610104 | 0.02939903 | 5.709821199 | 0.107750036 | 0.018871 |
| LOC111948210 | 5.6404303 | 5.76834671 | 0 | 1.13757403 | 5.704388504 | 0.568787015 | 0.099710427 |
| LOC110017180 | 5.36370416 | 6.03387933 | 0.55637983 | 3.86730384 | 5.698791742 | 2.211841831 | 0.388124699 |
| LOC110016418 | 4.8620656 | 6.51016387 | 0 | 0.06570976 | 5.686114735 | 0.032854879 | 0.005778089 |
| LOC110015392 | 4.89478152 | 6.43601282 | 0 | 0 | 5.665397169 | 0 | 0 |
| LOC105354559 | 5.16134815 | 6.15813299 | 1.11539444 | 2.11443318 | 5.659740574 | 1.61491381 | 0.285333539 |
| LOC110016132 | 5.72428931 | 5.51958709 | 0.67860973 | 0.21440471 | 5.621938202 | 0.446507218 | 0.079422292 |
| LOC111946906 | 5.33060982 | 5.87084613 | 0.2126719 | 0.26877219 | 5.600727975 | 0.240722046 | 0.042980492 |
| LOC110016651 | 4.35464201 | 6.78613134 | 1.29059795 | 1.08736148 | 5.570386671 | 1.188979714 | 0.213446531 |
| LOC111947082 | 2.72476171 | 8.35966554 | 0 | 0 | 5.542213626 | 0 | 0 |
| LOC110016391 | 5.63515196 | 5.42395169 | 0 | 0 | 5.529551824 | 0 | 0 |
| LOC105355673 | 5.30595064 | 5.7454746 | 6.31328585 | 6.23971734 | 5.525712621 | 6.276501597 | 1.135871883 |
| LOC111946823 | 4.04359615 | 6.99819794 | 1.29059795 | 1.42716194 | 5.520897044 | 1.358879945 | 0.246133904 |
| LOC105356549 | 4.98128284 | 6.04942281 | 0.25835553 | 0.89789321 | 5.515352823 | 0.578124372 | 0.104820923 |
| LOC111947470 | 5.23677927 | 5.77394317 | 0 | 0.10726674 | 5.505361217 | 0.05363337 | 0.009742026 |
| LOC110017190 | 5.63130593 | 5.35487399 | 2.66449048 | 2.07221607 | 5.493089959 | 2.368353273 | 0.431151372 |
| LOC111947184 | 4.3903512 | 6.59456664 | 0.85390016 | 0.17985813 | 5.492458919 | 0.516879143 | 0.094107057 |
| LOC110014683 | 4.05815574 | 6.81816693 | 0 | 0 | 5.438161336 | 0 | 0 |
| LOC105356387 | 5.69219049 | 5.13191856 | 0.1553826 | 0.09818533 | 5.412054525 | 0.126783968 | 0.023426218 |
| LOC110015956 | 4.61554262 | 6.1851108 | 0.33018803 | 2.08643835 | 5.400326708 | 1.208313188 | 0.223748164 |
| LOC105357135 | 4.46791568 | 6.274182 | 0.41503811 | 0 | 5.371048841 | 0.207519054 | 0.038636598 |
| LOC110016868 | 4.95411221 | 5.69977196 | 0 | 4.87088516 | 5.326942082 | 2.435442581 | 0.457193366 |
| LOC111948199 | 4.60973087 | 5.97452367 | 0.56812252 | 0.0598322 | 5.29212727 | 0.313977359 | 0.05932914 |
| LOC105356053 | 5.21879764 | 5.33715206 | 0.28492031 | 0.18003944 | 5.277974847 | 0.232479876 | 0.044047174 |
| LOC110016045 | 4.85446052 | 5.70122784 | 0.0974624 | 0.22581498 | 5.277844181 | 0.16163869 | 0.030625893 |
| LOC110015978 | 4.59642664 | 5.82882662 | 0.76286357 | 0 | 5.212626633 | 0.381431783 | 0.073174584 |
| LOC110013749 | 5.06174248 | 5.35201089 | 0 | 0 | 5.206876685 | 0 | 0 |
| LOC105356641 | 4.66568787 | 5.72579831 | 0 | 0 | 5.19574309 | 0 | 0 |
| LOC110017495 | 4.55230541 | 5.7559463 | 0.17171382 | 0.43401974 | 5.154125857 | 0.302866781 | 0.058762007 |
| LOC110017724 | 5.20510959 | 5.09171212 | 1.87801297 | 0.89002885 | 5.148410854 | 1.384020908 | 0.268824876 |
| LOC105355361 | 4.82034195 | 5.44857158 | 8.42133188 | 4.98880231 | 5.134456765 | 6.705067093 | 1.305896105 |
| LOC110017288 | 4.65282857 | 5.59314551 | 0 | 0 | 5.122987038 | 0 | 0 |
| LOC110016716 | 5.48275223 | 4.75753323 | 0.68936817 | 1.08901904 | 5.12014273 | 0.889193607 | 0.173665785 |
| LOC105355834 | 4.86564592 | 5.34458335 | 0.56079554 | 0 | 5.105114632 | 0.280397769 | 0.054924872 |
| LOC105354858 | 4.49717953 | 5.68132609 | 2.7440869 | 3.46794413 | 5.08925281 | 3.106015516 | 0.610308749 |
| LOC110015522 | 4.04196764 | 6.07887256 | 1.97306074 | 7.48059152 | 5.060420096 | 4.726826132 | 0.934077812 |
| LOC110015242 | 4.91350473 | 5.17720634 | 0 | 0.58557089 | 5.045355536 | 0.292785447 | 0.058030687 |
| LOC110014867 | 5.20161659 | 4.86685104 | 5.74006806 | 5.80338335 | 5.034233819 | 5.771725704 | 1.146495358 |
| LOC110016551 | 4.9938359 | 5.0671894 | 0.32375825 | 0.15343567 | 5.030512651 | 0.238596961 | 0.04742995 |
| LOC105356928 | 4.58585422 | 5.47149685 | 4.36051958 | 6.26224133 | 5.028675531 | 5.311380455 | 1.056218565 |
| LOC111949249 | 5.09204134 | 4.94183127 | 0.21559188 | 0.34057804 | 5.016936304 | 0.278084957 | 0.055429238 |
| LOC105353672 | 5.43035746 | 4.57347861 | 3.31349298 | 4.1875527 | 5.001918032 | 3.75052284 | 0.749816933 |
| LOC105355823 | 4.98020264 | 5.00682157 | 7.00473237 | 6.19674775 | 4.993512105 | 6.60074006 | 1.321863234 |
| LOC110016363 | 6.97715263 | 3.00437216 | 0 | 2.40699626 | 4.990762398 | 1.203498131 | 0.241145147 |
| LOC105356522 | 4.19334355 | 5.78007431 | 0.09456037 | 0 | 4.986708932 | 0.047280186 | 0.00948124 |
| LOC101170462 | 4.84308496 | 5.06059094 | 0.65527268 | 0.96614672 | 4.951837949 | 0.810709703 | 0.163718949 |
| LOC105357904 | 5.02072626 | 4.85704339 | 0 | 0 | 4.938884827 | 0 | 0 |
| LOC110015671 | 4.8416299 | 5.0202006 | 6.69633053 | 4.05506399 | 4.930915245 | 5.37569726 | 1.09020273 |
| LOC111948458 | 4.0849345 | 5.75827853 | 1.14522266 | 3.61829665 | 4.921606514 | 2.381759652 | 0.483939471 |
| LOC110017690 | 4.05336453 | 5.75734541 | 2.33587563 | 0 | 4.905354972 | 1.167937815 | 0.238094454 |
| LOC110015658 | 3.71220942 | 6.07423472 | 0 | 0 | 4.893222071 | 0 | 0 |
| LOC105353762 | 4.8225871 | 4.93195607 | 0 | 0 | 4.877271587 | 0 | 0 |
| LOC111946375 | 3.85873061 | 5.85571212 | 0.25823751 | 1.14225113 | 4.857221365 | 0.700244317 | 0.144165617 |
| LOC110016366 | 3.60214013 | 6.05200842 | 0 | 0.16864884 | 4.827074273 | 0.084324421 | 0.017469054 |
| LOC111946955 | 4.46714509 | 5.15208816 | 0.12247911 | 0 | 4.809616622 | 0.061239553 | 0.012732731 |
| LOC110015777 | 3.33814605 | 6.25872526 | 0 | 0.27350555 | 4.798435657 | 0.136752774 | 0.028499449 |
| LOC110016264 | 4.83367715 | 4.76021253 | 0.3714073 | 0.07823001 | 4.79694484 | 0.224818655 | 0.04686705 |
| LOC110017523 | 3.60100667 | 5.98434207 | 0.31127858 | 0 | 4.79267437 | 0.155639291 | 0.032474414 |
| LOC110016333 | 3.93894991 | 5.6339571 | 0.11429072 | 0.18054905 | 4.786453502 | 0.147419886 | 0.030799398 |
| LOC110017262 | 2.76345001 | 6.78269009 | 0 | 0.36227003 | 4.773070053 | 0.181135013 | 0.037949372 |
| LOC105354261 | 3.51582157 | 5.99259178 | 3.64698002 | 1.92042067 | 4.754206675 | 2.783700344 | 0.585523629 |
| LOC110016254 | 3.73468196 | 5.72906833 | 0 | 0.50999178 | 4.731875145 | 0.254995892 | 0.053888973 |
| LOC110016815 | 5.10892821 | 4.35399247 | 10.5990357 | 4.46497807 | 4.73146034 | 7.532006868 | 1.591898976 |
| LOC110015847 | 4.79005534 | 4.67084065 | 0 | 0 | 4.730447997 | 0 | 0 |
| LOC105358280 | 4.04267317 | 5.37466133 | 0.62902289 | 1.98737896 | 4.708667245 | 1.308200925 | 0.277828281 |
| LOC110016802 | 3.32867923 | 6.07887256 | 3.4528563 | 2.80522182 | 4.703775893 | 3.129039059 | 0.665218567 |
| LOC105356895 | 3.85397696 | 5.51793105 | 2.39865022 | 6.56800168 | 4.685954006 | 4.483325951 | 0.95675842 |
| LOC101170150 | 3.98146899 | 5.35758247 | 1.3663153 | 2.6685872 | 4.669525732 | 2.017451249 | 0.432046286 |
| LOC110014979 | 3.9023546 | 5.39939083 | 0 | 0 | 4.650872715 | 0 | 0 |
| LOC105358717 | 4.64939499 | 4.53368105 | 0.67295465 | 0 | 4.591538016 | 0.336477323 | 0.073282051 |
| LOC105358373 | 3.46941709 | 5.71157273 | 0 | 0 | 4.590494908 | 0 | 0 |
| LOC110014922 | 4.39477696 | 4.77534658 | 2.84920314 | 3.24070989 | 4.585061767 | 3.044956512 | 0.664103706 |
| LOC111947933 | 3.26168529 | 5.90154412 | 2.08206962 | 0.32891183 | 4.581614705 | 1.205490725 | 0.263114819 |
| LOC101167119 | 4.90212002 | 4.1777439 | 0.42374955 | 0 | 4.53993196 | 0.211874776 | 0.046669152 |
| LOC105355032 | 4.43521015 | 4.63439821 | 2.40034778 | 1.13757403 | 4.53480418 | 1.768960905 | 0.390085401 |
| LOC111947329 | 4.28935502 | 4.59551829 | 0 | 0 | 4.442436654 | 0 | 0 |
| LOC110015119 | 3.72131808 | 5.16028737 | 1.30852292 | 1.32295646 | 4.440802725 | 1.315739693 | 0.296284202 |
| LOC111947720 | 3.76899505 | 5.1015046 | 0 | 0.43596206 | 4.435249823 | 0.217981029 | 0.049147407 |
| LOC105354785 | 3.15673613 | 5.69204869 | 0 | 0.21780381 | 4.424392411 | 0.108901904 | 0.024613979 |
| LOC105354668 | 4.77456069 | 4.06397904 | 0.61523934 | 0.07775321 | 4.419269862 | 0.346496273 | 0.078405774 |
| LOC110015667 | 3.08829857 | 5.68500571 | 0.73223044 | 0 | 4.386652142 | 0.366115222 | 0.083461193 |
| LOC111948779 | 3.60418216 | 5.16028737 | 0 | 3.77987561 | 4.382234765 | 1.889937806 | 0.431272606 |
| LOC101157740 | 3.86058707 | 4.88322496 | 0.28102506 | 0.79910122 | 4.371906013 | 0.540063139 | 0.123530364 |
| LOC110016393 | 4.10557577 | 4.62712115 | 0 | 0.10983956 | 4.366348457 | 0.05491978 | 0.012577965 |
| LOC105355424 | 3.77856016 | 4.93765556 | 0.16331334 | 0.03439891 | 4.358107857 | 0.098856123 | 0.022683267 |
| LOC110015648 | 2.96997504 | 5.68525791 | 0.4739647 | 0.09983182 | 4.327616474 | 0.286898261 | 0.066294752 |
| LOC110015169 | 3.98661065 | 4.60880628 | 4.67472311 | 4.31728159 | 4.297708463 | 4.496002351 | 1.046139446 |
| LOC105357301 | 3.13420951 | 5.45581043 | 0 | 0 | 4.295009971 | 0 | 0 |
| LOC110015175 | 4.93728057 | 3.62914888 | 0.64018336 | 1.11245207 | 4.283214721 | 0.876317711 | 0.204593458 |
| LOC111947354 | 4.37863453 | 4.1580828 | 0.06488543 | 0.2460043 | 4.268358662 | 0.155444868 | 0.036417949 |
| LOC105355864 | 4.04096017 | 4.46995837 | 0 | 0.2162217 | 4.255459269 | 0.108110849 | 0.025405213 |
| LOC110015267 | 3.89664016 | 4.57538192 | 3.89230123 | 3.78387972 | 4.236011039 | 3.838090475 | 0.906062435 |
| LOC110015168 | 4.35381366 | 4.08150559 | 0 | 0.71344523 | 4.217659623 | 0.356722615 | 0.084578332 |
| LOC111948948 | 3.7512364 | 4.62100828 | 3.360562 | 3.12939639 | 4.186122339 | 3.244979195 | 0.775175433 |
| LOC110016617 | 4.09738604 | 4.19030854 | 1.59383995 | 0 | 4.143847288 | 0.796919975 | 0.192314031 |
| LOC105356886 | 2.95414216 | 5.33141935 | 0 | 0 | 4.142780756 | 0 | 0 |
| LOC110017305 | 3.64747407 | 4.63753756 | 1.02257942 | 0.38769708 | 4.142505818 | 0.70513825 | 0.17022022 |
| LOC111947788 | 4.58110522 | 3.66651997 | 1.2396533 | 1.8277688 | 4.123812596 | 1.533711047 | 0.371915797 |
| LOC105356923 | 3.45963615 | 4.75812837 | 3.71244682 | 3.44061357 | 4.108882261 | 3.576530195 | 0.870438715 |
| LOC110016136 | 3.98474964 | 4.22986903 | 0.41857231 | 0 | 4.107309331 | 0.209286154 | 0.050954563 |
| LOC105354589 | 3.78059549 | 4.42532167 | 0.18899428 | 1.55251661 | 4.102958578 | 0.870755448 | 0.212226234 |
| LOC111949299 | 3.58371582 | 4.59577994 | 0.35403877 | 0.22371498 | 4.089747882 | 0.288876873 | 0.070634396 |
| LOC111947537 | 4.59918519 | 3.55374629 | 0 | 0.93787986 | 4.076465739 | 0.468939932 | 0.115035907 |
| LOC110015789 | 4.19474695 | 3.91684463 | 0 | 0 | 4.055795789 | 0 | 0 |
| LOC111948159 | 3.86621594 | 4.236317 | 1.33681531 | 0.60337541 | 4.05126647 | 0.970095362 | 0.239454839 |
| LOC111948408 | 3.75080964 | 4.31139368 | 0.12861932 | 0.52827954 | 4.031101659 | 0.328449431 | 0.081478826 |
| LOC105354344 | 4.51042898 | 3.54824514 | 0.83976514 | 1.21289727 | 4.029337059 | 1.026331201 | 0.254714656 |
| LOC110015779 | 2.87933663 | 5.11436713 | 1.25757932 | 0.99332104 | 3.996851877 | 1.125450177 | 0.28158416 |
| LOC111946474 | 5.33370581 | 2.64940929 | 1.58076595 | 1.39842714 | 3.991557551 | 1.489596541 | 0.373186788 |
| LOC110015793 | 3.28456601 | 4.67346781 | 0.29626934 | 0 | 3.979016906 | 0.148134671 | 0.037228962 |
| LOC110017519 | 5.42977878 | 2.52405361 | 0 | 0 | 3.976916191 | 0 | 0 |
| LOC111947793 | 4.0855712 | 3.85682378 | 3.58598669 | 3.91393694 | 3.971197487 | 3.749961818 | 0.944289935 |
| LOC105358128 | 2.87826942 | 5.05685799 | 0.30621988 | 0 | 3.967563703 | 0.153109941 | 0.038590418 |
| LOC111948913 | 3.53209852 | 4.38624426 | 0 | 0 | 3.959171391 | 0 | 0 |
| LOC105354265 | 3.76493424 | 4.11894403 | 0.54493757 | 0.68868556 | 3.941939137 | 0.616811565 | 0.156474147 |
| LOC105357535 | 5.06932412 | 2.80815638 | 0 | 0 | 3.93874025 | 0 | 0 |
| LOC110015181 | 3.98745617 | 3.87399135 | 0.13787364 | 0.52272914 | 3.930723756 | 0.330301387 | 0.084030679 |
| LOC110015111 | 3.36727399 | 4.46742313 | 0 | 0 | 3.917348564 | 0 | 0 |
| LOC105354494 | 3.48078911 | 4.27167376 | 0 | 0 | 3.876231431 | 0 | 0 |
| LOC110016934 | 3.39726595 | 4.34288849 | 0.16018189 | 0.70852585 | 3.870077221 | 0.434353871 | 0.112233903 |
| LOC110014286 | 3.44393674 | 4.24716589 | 0.42028394 | 0 | 3.845551313 | 0.210141971 | 0.054645473 |
| LOC101157968 | 4.57302665 | 3.11782398 | 0.13176734 | 0.99915593 | 3.845425314 | 0.565461637 | 0.147047879 |
| LOC111948471 | 3.39866928 | 4.27020216 | 1.00727353 | 0.25459604 | 3.834435721 | 0.630934783 | 0.164544363 |
| LOC111947499 | 3.70121783 | 3.95527521 | 1.03531484 | 0.81776155 | 3.828246522 | 0.926538195 | 0.242026784 |
| LOC111947645 | 3.07094046 | 4.56812325 | 0 | 4.0257999 | 3.81953185 | 2.012899948 | 0.527001744 |
| LOC110015109 | 3.7740117 | 3.85960056 | 0.46512499 | 0 | 3.816806131 | 0.232562494 | 0.060931178 |
| LOC110013497 | 3.26910585 | 4.35399247 | 0 | 0.59798813 | 3.811549159 | 0.298994067 | 0.078444237 |
| LOC110017746 | 2.93227668 | 4.69039365 | 0.15598287 | 0 | 3.811335165 | 0.077991433 | 0.020463021 |
| LOC105356055 | 3.26971406 | 4.33464139 | 0.5024728 | 0.79377388 | 3.802177723 | 0.64812334 | 0.170461085 |
| LOC101171442 | 3.47955818 | 4.1059261 | 2.77643371 | 6.66676489 | 3.79274214 | 4.721599298 | 1.244903852 |
| LOC110016119 | 4.69786502 | 2.88264329 | 2.59899725 | 1.23171809 | 3.790254156 | 1.91535767 | 0.505337529 |
| LOC111947437 | 3.59680291 | 3.97864829 | 0 | 0.09622798 | 3.787725599 | 0.048113988 | 0.012702607 |
| LOC111948639 | 3.57345798 | 3.99710784 | 0 | 0.14639272 | 3.785282912 | 0.073196362 | 0.019337091 |
| LOC110015391 | 3.12393063 | 4.43718978 | 0 | 0 | 3.780560201 | 0 | 0 |
| LOC110017596 | 3.67714131 | 3.82320886 | 0.63571964 | 0.48204891 | 3.750175087 | 0.558884276 | 0.149028849 |
| LOC110014665 | 3.1772593 | 4.26695725 | 0.10865234 | 0.02288559 | 3.722108277 | 0.065768963 | 0.017669815 |
| LOC105356242 | 4.11625756 | 3.28554774 | 2.08283678 | 1.18451887 | 3.700902653 | 1.633677825 | 0.441426857 |
| LOC111947748 | 0 | 7.38859328 | 0 | 0 | 3.69429664 | 0 | 0 |
| LOC111948037 | 3.05740767 | 4.31490816 | 0.50743438 | 0.64128949 | 3.686157911 | 0.574361936 | 0.155815879 |
| LOC110016963 | 2.97246732 | 4.35715901 | 1.02778528 | 0.77934163 | 3.664813166 | 0.903563452 | 0.246551028 |
| LOC110017261 | 4.59773291 | 2.72221199 | 0 | 0 | 3.659972448 | 0 | 0 |
| LOC105355790 | 3.38479716 | 3.89425413 | 1.46294488 | 0.73954088 | 3.639525646 | 1.101242881 | 0.302578684 |
| LOC110016211 | 2.96603745 | 4.28745668 | 0.05916704 | 0 | 3.626747067 | 0.02958352 | 0.00815704 |
| LOC105356934 | 3.83594973 | 3.41675703 | 6.67453654 | 6.16418407 | 3.626353382 | 6.419360304 | 1.770197118 |
| LOC111947644 | 3.25885122 | 3.98079311 | 1.69021167 | 0.83069359 | 3.619822166 | 1.260452631 | 0.34820844 |
| LOC105354940 | 3.88667211 | 3.34721343 | 0 | 0 | 3.616942771 | 0 | 0 |
| LOC111948440 | 4.03071259 | 3.19464536 | 0.20905396 | 0.39629983 | 3.612678977 | 0.302676893 | 0.08378184 |
| LOC105357782 | 2.36707099 | 4.84150514 | 0 | 0 | 3.604288062 | 0 | 0 |
| LOC111947728 | 2.12872009 | 5.07965788 | 2.94417658 | 1.86040753 | 3.604188984 | 2.402292051 | 0.666527771 |
| LOC105356964 | 4.54848935 | 2.65808125 | 0 | 0 | 3.603285299 | 0 | 0 |
| LOC110015912 | 3.79800419 | 3.38295818 | 0.25417352 | 0.16061072 | 3.590481181 | 0.20739212 | 0.057761651 |
| LOC110014998 | 3.59929252 | 3.58143472 | 0.40362863 | 0.25505051 | 3.590363617 | 0.329339574 | 0.091728752 |
| LOC111948960 | 2.92245736 | 4.21938955 | 0.5349671 | 0.28170209 | 3.570923452 | 0.408334593 | 0.114349859 |
| LOC110017112 | 3.22457008 | 3.91601689 | 1.25432375 | 0.26419989 | 3.570293485 | 0.759261817 | 0.212660898 |
| LOC111947318 | 4.37202966 | 2.74040106 | 0.58517795 | 0.36977044 | 3.55621536 | 0.477474197 | 0.134264702 |
| LOC101156560 | 3.58521278 | 3.52549997 | 0 | 0 | 3.555356377 | 0 | 0 |
| LOC110015451 | 2.91107021 | 4.16792441 | 7.85113753 | 11.0670397 | 3.53949731 | 9.459088591 | 2.672438418 |
| LOC111948449 | 1.83566255 | 5.22377069 | 0 | 0.10462749 | 3.529716621 | 0.052313744 | 0.014820947 |
| LOC110016025 | 3.78439127 | 3.27479775 | 1.81180097 | 0.38162206 | 3.529594511 | 1.096711513 | 0.31071884 |
| LOC110014939 | 3.3304956 | 3.72534115 | 0 | 1.22795424 | 3.527918375 | 0.613977122 | 0.174033823 |
| LOC110015500 | 2.76659807 | 4.27851464 | 0.13999057 | 0 | 3.522556357 | 0.069995283 | 0.019870593 |
| LOC110017383 | 2.83829345 | 4.19273349 | 0 | 0 | 3.51551347 | 0 | 0 |
| LOC110013502 | 3.38479716 | 3.63463719 | 0 | 0 | 3.509717175 | 0 | 0 |
| LOC110017617 | 3.96328976 | 3.03987838 | 7.19449694 | 1.29890271 | 3.501584071 | 4.246699825 | 1.212793907 |
| LOC105356293 | 3.67242703 | 3.32823268 | 0.68136105 | 0.19570362 | 3.500329856 | 0.438532334 | 0.125283145 |
| LOC110016718 | 3.57345798 | 3.42609243 | 1.85338329 | 3.22063992 | 3.499775209 | 2.537011601 | 0.724907015 |
| LOC111948626 | 2.5867991 | 4.4091063 | 0.89443339 | 1.13037419 | 3.497952696 | 1.012403792 | 0.289427525 |
| LOC111947685 | 2.41842756 | 4.53433535 | 0 | 0 | 3.476381454 | 0 | 0 |
| LOC110013437 | 3.22942831 | 3.69450868 | 0 | 0.14351075 | 3.461968491 | 0.071755373 | 0.020726755 |
| LOC105354017 | 2.48802259 | 4.39495421 | 0.15641447 | 0.2965121 | 3.441488403 | 0.226463287 | 0.065803879 |
| LOC105355305 | 2.59501116 | 4.22959268 | 1.17767063 | 1.59463502 | 3.412301919 | 1.386152827 | 0.40622221 |
| LOC110014721 | 3.83769255 | 2.9435442 | 0 | 0 | 3.390618379 | 0 | 0 |
| LOC111946919 | 3.13539767 | 3.62024615 | 0.20982996 | 0.26518058 | 3.377821908 | 0.23750527 | 0.070313142 |
| LOC110016555 | 2.72274487 | 4.02204486 | 0.62762609 | 0.79318633 | 3.372394861 | 0.710406214 | 0.210653332 |
| LOC110013952 | 3.13975548 | 3.55997781 | 0.56641473 | 0.17895704 | 3.349866645 | 0.372685885 | 0.111253947 |
| LOC105354623 | 3.62546257 | 3.07042463 | 0.82265558 | 0.44556954 | 3.347943598 | 0.634112556 | 0.189403596 |
| LOC110017298 | 3.14728468 | 3.54052636 | 1.79558213 | 0.82517643 | 3.343905521 | 1.31037928 | 0.391870904 |
| LOC110015482 | 2.46957255 | 4.20929785 | 3.13096723 | 1.43886504 | 3.339435202 | 2.284916135 | 0.68422233 |
| LOC110017486 | 3.24376394 | 3.40947558 | 0 | 0 | 3.326619764 | 0 | 0 |
| LOC105353753 | 2.18330266 | 4.4656333 | 0 | 0 | 3.324467978 | 0 | 0 |
| LOC111947922 | 2.11690699 | 4.48446727 | 0.6273939 | 0.19822322 | 3.300687131 | 0.41280856 | 0.125067461 |
| LOC111946706 | 2.58627107 | 4.01004573 | 1.03848482 | 0.9843185 | 3.298158398 | 1.011401657 | 0.306656484 |
| LOC105355335 | 4.50373837 | 2.07264435 | 0 | 0 | 3.288191359 | 0 | 0 |
| LOC111948475 | 3.13706118 | 3.43736248 | 0 | 0 | 3.287211829 | 0 | 0 |
| LOC110015760 | 3.14005997 | 3.41845059 | 0.42028394 | 0.2655749 | 3.27925528 | 0.342929423 | 0.104575397 |
| LOC105357086 | 2.75708617 | 3.69466493 | 0.59171169 | 0 | 3.22587555 | 0.295855846 | 0.091713348 |
| LOC105356905 | 3.02923908 | 3.41567482 | 0 | 0 | 3.222456949 | 0 | 0 |
| LOC111946473 | 4.08922891 | 2.30434885 | 1.2119367 | 1.03932108 | 3.196788879 | 1.125628889 | 0.352112364 |
| LOC110017679 | 2.75065019 | 3.64039275 | 0 | 0 | 3.195521469 | 0 | 0 |
| LOC110015152 | 2.52292751 | 3.80231701 | 0.27547851 | 1.39258575 | 3.162622261 | 0.834032131 | 0.26371538 |
| LOC111947548 | 2.3613959 | 3.94277351 | 0.19995823 | 0 | 3.152084704 | 0.099979113 | 0.031718409 |
| LOC105355793 | 2.7047267 | 3.58564086 | 0 | 0.26264577 | 3.145183778 | 0.131322885 | 0.041753644 |
| LOC110014740 | 3.20089482 | 3.06889337 | 0 | 1.04904037 | 3.134894093 | 0.524520184 | 0.167316716 |
| LOC111946566 | 1.53306173 | 4.70348774 | 0 | 0 | 3.118274733 | 0 | 0 |
| LOC110014159 | 2.71390609 | 3.51557426 | 0 | 0 | 3.114740177 | 0 | 0 |
| LOC111948340 | 2.81983028 | 3.378142 | 0.83572132 | 0.42246984 | 3.098986138 | 0.629095579 | 0.203000449 |
| LOC111946404 | 4.06525275 | 2.13202386 | 0.21625168 | 0 | 3.098638302 | 0.108125842 | 0.034894632 |
| LOC110016492 | 2.50118629 | 3.65415629 | 0 | 0.26766448 | 3.077671286 | 0.133832239 | [illegible] |
| [illegible] | [illegible] | [illegible] | [illegible] | [illegible] | [illegible] | [illegible] | [illegible] |

| | | | | | | | |
|---|---|---|---|---|---|---|---|
| LOC105357759 | 1.45355863 | 2.70276933 | 0.8224276 | 0 | 2.078163978 | 0.411213799 | 0.197873605 |
| LOC110015792 | 1.47714355 | 2.67267594 | 0 | 0 | 2.074909746 | 0 | 0 |
| LOC110015264 | 1.78294646 | 2.35966951 | 0 | 2.47481463 | 2.071307988 | 1.237407316 | 0.597403826 |
| LOC105354622 | 2.04357129 | 2.08991639 | 4.23961427 | 13.3949342 | 2.066743835 | 8.817274234 | 4.266263716 |
| LOC111949246 | 1.5250532 | 2.59939849 | 0 | 0 | 2.062225843 | 0 | 0 |
| LOC111946421 | 2.16939627 | 1.9523635 | 0 | 0.1137574 | 2.060879885 | 0.056878702 | 0.027599232 |
| LOC110013763 | 1.16442808 | 2.95503636 | 0 | 0 | 2.05973222 | 0 | 0 |
| LOC110015172 | 1.93795285 | 2.18009296 | 0 | 0 | 2.059022906 | 0 | 0 |
| LOC111948748 | 2.13733839 | 1.97427999 | 0.28607384 | 0.45192086 | 2.055809187 | 0.368997347 | 0.179490076 |
| LOC111946962 | 2.01834201 | 2.06411495 | 0 | 0 | 2.041228479 | 0 | 0 |
| LOC110015353 | 1.76805014 | 2.31040985 | 0 | 1.03013193 | 2.039229992 | 0.515065963 | 0.252578652 |
| LOC110017289 | 2.02640021 | 2.00691308 | 0 | 0.02796291 | 2.016656643 | 0.013981457 | 0.006932988 |
| LOC101165136 | 1.69872925 | 2.31633847 | 0 | 0 | 2.007533859 | 0 | 0 |
